# Stalled translation on transcripts cleaved by RNase L activates signaling important for innate immunity

**DOI:** 10.1101/2025.06.10.658914

**Authors:** Agnes Karasik, Grant D. Jones, Nicholas R. Guydosh

**Affiliations:** Laboratory of Biochemistry and Genetics, National Institute of Diabetes and Digestive and Kidney Diseases, National Institutes of Health, Bethesda, MD 20892

**Keywords:** endonuclease, mRNA fragments, PELO, nanopore, ribosome profiling

## Abstract

RNase L is an endonuclease that responds to infections by cleaving most host- and pathogen-derived single-stranded RNAs. This widespread RNA cleavage can lead to death of the infected cell via the ribotoxic stress response (RSR). An ongoing challenge is to understand how RNase L’s endonuclease activity triggers cell death to benefit the host. To address this question, we used nanopore-based long-read sequencing to show that 3’ mRNA fragments in the cell were not fully degraded after RNase L activation and that these fragments were translated by ribosomes. We further asked whether ribosomes on mRNA fragments stall when they reach 3’ ends created by RNase L. We used ribosome profiling to capture footprints protected by these ribosomes, which can be identified by their short length (15-18 nt). We found that RNase L activation increased the number of stalled ribosomes at RNase L cleavage sites. Loss of the ribosome rescue factor PELO increased the number of short footprints derived from stalled ribosomes and augmented the RSR. Our work therefore establishes a role for fragmented mRNA in causing ribosome stalling that promotes innate immunity via the RSR.

**Graphical Abstract:** Stalled translation of mRNAs that are fragmented by RNase L leads to ribosome stalling and potentially collisions. Stalled ribosomes are rescued by PELO or activate innate immune signaling via ZAK⍺. Renderings based on PDB 4o1o and 3jag.

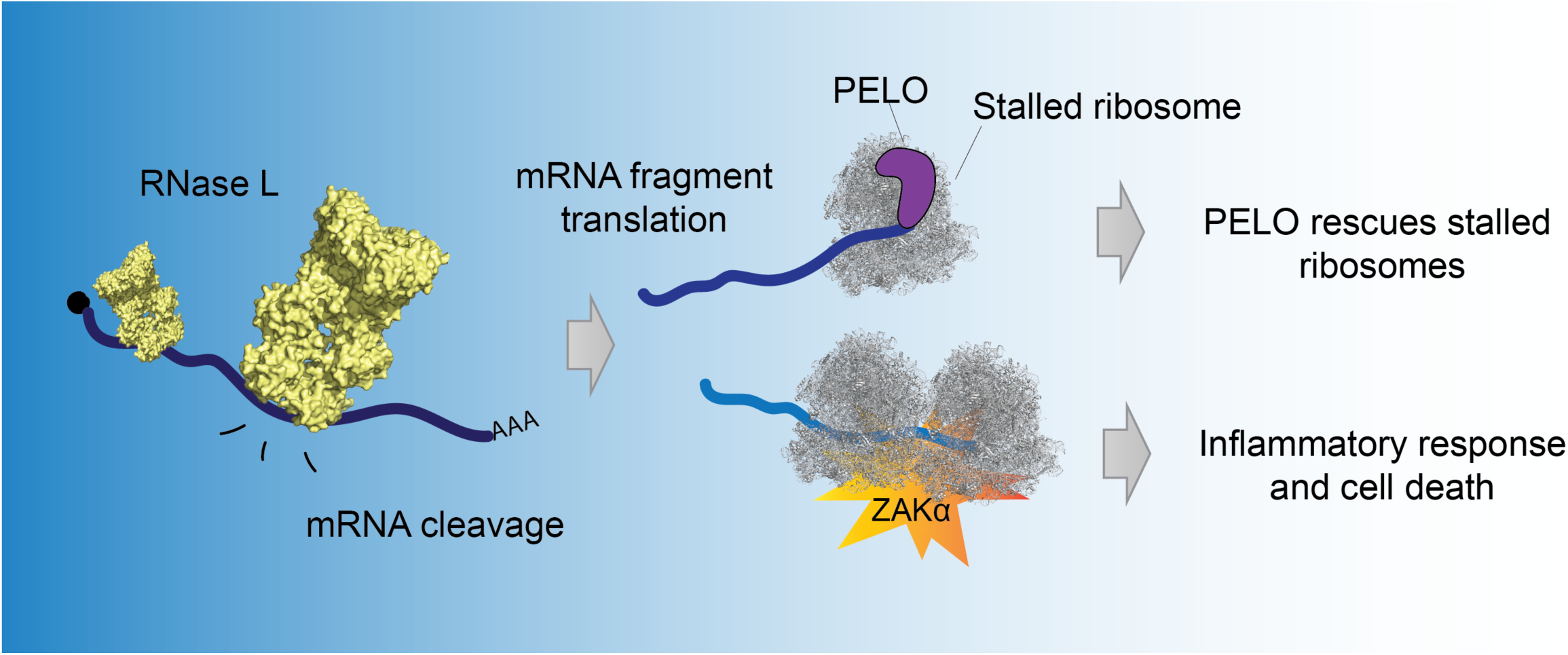

**Highlights:**

- Activation of RNase L leads to accumulation and translation of mRNA fragments
- Ribosomes stall at the 3’ end of the RNase L cleaved mRNA fragments
- PELO rescues ribosomes stalled due to RNase L activation

## Introduction

Endonucleolytic cleavage of messenger RNAs (mRNAs) plays an important role in in cell homeostasis by tuning transcript abundance and thus altering gene expression ^1^. When particularly widespread, endonucleolytic cleavage of mRNAs leads to inflammation and cell death ^2,3^. An endonuclease known to have this role, Ribonuclease L (RNase L), is an important factor in the innate immune response, particularly in humans where it is linked to many diseases ^4–7^. One common signature of viral infection is the accumulation of viral dsRNA in the cytoplasm. This dsRNA is sensed by cytoplasmic dsRNA-binding proteins, including <u>O</u>ligo<u>A</u>denylate <u>S</u>ynthetases (OAS1-3). When OAS binds to dsRNA, it produces a small molecule, 2’-5’ oligoadenylate (2-5A), a potent and specific activator of RNase L. Active RNase L cleaves viral single-stranded RNA genomes, RNA replication intermediates, and mRNAs, and therefore promotes viral elimination. RNase L activation also induces other processes in the cell and can ultimately cause cell death and benefit the host organism by eliminating the infected cell^3,8,9^. In particular, RNase L widely cleaves host-encoded RNA, including mRNA, and therefore causes its degradation ^10,11^. We and others found that this endonucleolytic activity of RNase L also leads to activation of the MAP3 kinase ZAKα^12,13^. ZAKα senses ribosome stalling and collisions (queuing behind a ribosome that stalls) that can be caused by cellular stress, such as UV radiation ^14,15^. ZAKα activation triggers further downstream signaling that is known as the Ribotoxic Stress Response (RSR) ^16^. This includes activation of the JNK and p38 MAPKs that induce an inflammatory response and ultimately cell death. However, the exact molecular mechanism of how RNase L activation causes ribosome stalling (and likely collisions) that leads to ZAKα activation is unclear.

Several hints for how mRNA fragmentation that is caused by RNase L would lead to ribosome stalling have come from prior studies. In particular, ribosomes stall and collide upon encountering endonuclease-cleaved 3’ ends of mRNAs fragments formed during the <u>U</u>nfolded<u> P</u>rotein <u>R</u>esponse (UPR) in yeast, suggesting a model that points to translation of the fragmented mRNA as being critical. These stalling events were found to be rescued (ribosomes were removed from the mRNA) by Dom34, the yeast ortholog of the human PELO (Protein pelota homolog), that forms a complex with HBS1L, a GTPase ^17^. The PELO-HBS1L complex recognizes stalled ribosomes, leading to recruitment of ABCE1, a recycling factor ^18^ that splits the stalled 80S ribosome so it can dissociate from the mRNA. Whether the PELO-HBS1L complex has a role in rescuing stalled ribosomes after RNase L activation remains unknown.

In support of the possibility that mRNA fragments generated by RNase L are translated, we previously reported the observation of ribosome footprints within open reading frames (ORFs) in non-coding regions of mRNAs when RNase L was active using ribosome profiling^19^. Since RNase activity was necessary to cause this effect, we hypothesized that ribosomes could initiate translation on mRNA fragments that are generated by RNase L and translate nearby ORFs in noncoding regions ^19^. These elongating ribosomes would be expected to stall if they encountered the cleaved 3’ ends or poly(A) tail of mRNA fragments. These stalled, and potentially collided, ribosomes would provide an interface for ZAKα activation.

Several methodologies can be used to ask the question of whether mRNA fragments are present in the cell. Earlier studies used sequencing methodologies such as RtcB-seq and short-read Illumina sequencing to suggest that RNA fragments (including mRNA fragments) may exist for some time in the cell when RNase L is active ^19–21^. However, these methodologies are limited in their ability to determine if a read originates from fragmented vs full-length mRNA. Thus, it remains unclear how long mRNA fragments survive decay in the cell during the innate immune response and whether they could be translated or serve other functions. Recent technological advances in nanopore technologies that sequence the full length of mRNA ^22^ allow for the study of endonuclease cleavage products in cells in a high-throughput manner. For instance, in a recent study using nanopore sequencing, shortening of poly(A)-tailed mRNAs was observed and attributed to an unidentified endonuclease during arsenite stress ^23^.

Several factors could affect the stability of mRNA fragments. In general, mRNAs are protected from degradation by their 5’ end cap structure and 3’ end poly(A) tail. When endonucleolytic cleavage occurs, the resulting fragments lack one of these features and are therefore susceptible to RNA degradation by exonucleases. The exosome eliminates RNA fragments lacking a poly(A) tail in 3’ to 5’ direction while XRN1 degrades uncapped RNAs with a 5’ phosphate in the 5’ to 3’ direction. These decay mechanisms are thought to eliminate unprotected mRNA fragments rapidly, observed within minutes ^24,25^ Under certain circumstances, such as RNase L activation, a large amount of RNA fragments could overwhelm these decay machineries, making RNA fragments have a longer half-life^19^. The 5’ fragments are characterized by a 2’-3’-cyclic phosphate at their 3’ end and the 3’ fragments by a 5’ OH terminus ^26,27^. During RNase L activation, the exosome has a role in degrading 5’ fragments and loss of the exosome was shown to enhance some downstream effects of RNase L activation, such as activation of the interferon response ^28^. In contrast, 3’ fragments are mostly degraded by XRN1^29^ and it is assumed that this enzyme is responsible for elimination of RNase L cleaved fragments during the innate immune response. Because the 3’ fragments lack a 5’ phosphate, which is the preferred substrate for XRN1, the XRN1-mediated decay process may occur inefficiently or an additional protein is involved in phosphorylating the 5’ end ^30–33^.

In the present study, we asked how active RNase L induces downstream signaling via ribosome stalling. We present evidence that mRNA fragments created by RNase L are detectable in the cell and translated by utilizing direct RNA nanopore sequencing. Additionally, we show that stalled ribosomes are detectable at the end of cleaved mRNAs and that these are rescued by PELO, allowing it to attenuate downstream signaling via the RSR. Our work therefore reveals how endonucleolytic cleavage of mRNAs is linked via translation to physiological outcomes of the innate immune response.

## Results

### Activation of RNase L leads to accumulation of mRNA fragments

Previous work showed that poly(A)-selected RNA samples from permeabilized cells where RNase L had been activated and subjected to RNA-Seq (poly(A)+ “short-read” RNA-seq) had higher levels of reads that mapped to the 3’ ends of transcripts as compared to 5’ ends ^21^. The enrichment of these 3’ end mapped reads implies that mRNAs cleaved by RNase L are present in the cytoplasm. However, this approach has limitations that make it difficult to determine the nature and abundance of the cleaved mRNAs. For instance, mRNA is fragmented during RNA-Seq library preparation (hence the term “short read sequencing”). This fragmentation step during the library preparation protocol makes it challenging to identify mRNA fragments generated by RNase L. Also, during library preparation, the fragmented samples must be reverse transcribed and then amplified by Polymerase Chain Reaction (PCR), which can introduce bias ^34^.

To address the question of whether mRNA fragments generated by RNase L are abundant in cells, we adopted direct “long-read” RNA nanopore sequencing ^22,23^. This approach circumvents the disadvantages of traditional RNA-Seq by directly sequencing the RNA and generates full transcript-length reads. First, we transfected wild type (WT) and exonuclease deficient (*XRN1* KO) A549 human lung carcinoma cells with the RNase L activator, 2-5A. We reasoned that any 3’ end fragment of mRNA that is produced by RNase L cleavage would be stabilized in *XRN1* KO cells and thus improve the odds of detection. Active RNase L cleaves rRNA at specific sites^11^ and these cleavage products can be visualized by separating the RNA by molecular weight. Thus, as is standard in the field, we used total RNA (containing rRNA) from these samples to assess RNase L activation (Figure S1A, red arrows). We found that WT and *XRN1* KO cells exhibited similar levels of RNase L activation in 2-5A treated samples. Next, we prepared nanopore sequencing libraries and performed direct RNA sequencing (Figure 1A, see Methods for more details). To assess differences between RNase L activated (+2-5A) and control cells (transfection reagent treated but untreated by 2-5A), we first aligned reads to the human genome (see Methods for details). As the nanopores sequence the RNA from the 3’ to 5’ direction, we expected to observe an enrichment for reads that consisted of only a 3’ portion of the full-length RNA in 2-5A treated samples as compared to a control. These short reads that don’t span the entire transcript length are expected to increase in abundance due to mRNA fragmentation by RNase L. Since only poly(A)-tailed transcripts are selected for sequencing through nanopores, 5’ mRNA fragments are not sequenced in these experiments. It is known that a basal level of shorter reads exists in nanopore data and is thought to arise from pore clogging and degradation of the samples during preparation ^35,36^. Our analysis therefore focused on shortened reads in excess of this background level in control samples prepared at the same time.

**Figure 1.**
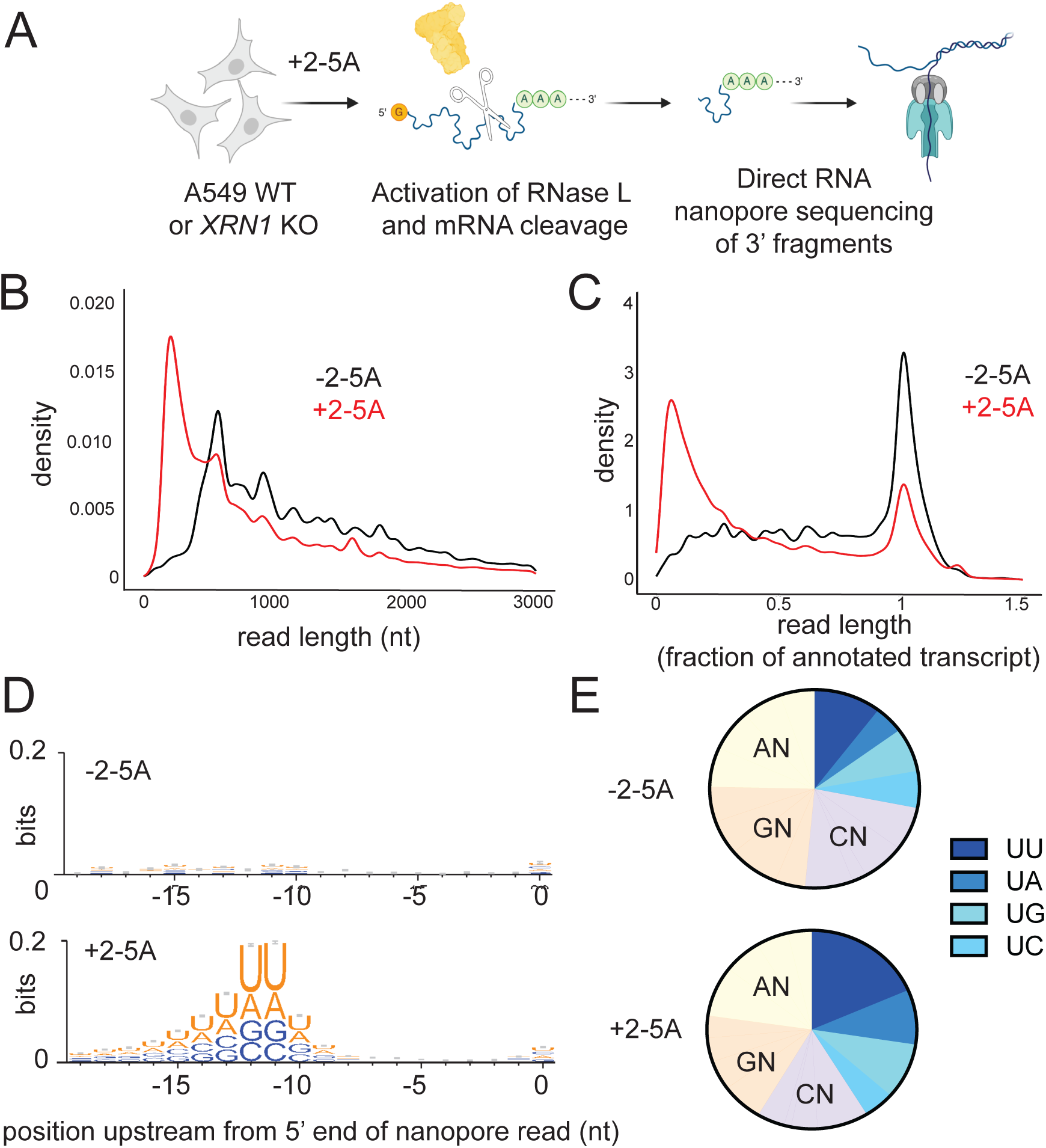
RNase L 3’ cleavage products are detectable after RNase L activation. **A** Schematic representation of direct RNA nanopore sequencing experiments. A549 cells were transfected with 5 µM of 2-5A for 4.5 hours then total RNA was extracted. RNase L 3’ cleavage products and uncleaved mRNA that contain poly(A) were sequenced. **B** Length distribution of nanopore direct sequencing reads in 2-5A treated (+2-5A) and control (−2-5A) samples in *XRN1* KO cells. **C** Normalized length distribution of nanopore direct sequencing reads to their respective annotated reference transcript in 2-5A treated (+2-5A) and control (−2-5A) samples in *XRN1* KO cells. **D** Weblogo analysis shows enrichment of U bases in transcriptome positions just upstream of where nanopore sequencing stopped in ±2-5A treated *XRN1* KO cells. **E** Di-nucleotide motif distribution near the 5’ end of 3’ fragments in ±2-5A treated *XRN1* KO cells (−11-−12 positions to account for RNA left in pore after sequencing stops). Number of total fragments were for were 212,139 (-2-5A) and 1,01262 x 10^6^ (+2-5A). In both D and E, 3’ fragments were defined as reads that were shorter than a third of their respective annotated transcript.

We observed short reads that spanned a 3’ portion of the transcript on individual genes in RNase L activated samples as compared to their respective control in both WT and *XRN1* KO cells (Figure S1B, *ACTG1* is shown as an example). This trend was stronger in the *XRN1* KO cells, supporting the interpretation that short reads derive from mRNAs that are cleaved and degraded by XRN1 after 2-5A treatment. Interestingly, we noted a small population of full-length reads in 2-5A treated cells. These full-length transcripts could arise from mRNAs that are protected from cleavage by RNase L or from technical artifacts, such as less than 100% efficiency of 2-5A transfection. Next, we assessed the global length distribution of transcriptome-mapped reads from the nanopore sequencing. We found that shorter reads were more abundant in the length distribution from RNase L activated samples as compared to control samples in both WT and *XRN1* KO cells (Figure 1B, S1C-D). As expected, reads from *XRN1* KO cells exhibited a more pronounced shift to shorter lengths as compared to WT, presumably due to stabilization of these 3’ end fragments during RNase L activation. We also normalized the detected read lengths to their corresponding reference transcript length (using MANE reference sequence, see Materials and Methods) to more clearly show the strong emergence of 3’ fragments of mRNA in the samples where RNase L was active, as compared to control samples, and that the lack of *XRN1* amplified this effect (Figure 1C, S1E-F).

To further validate that the detected 3’ mRNA fragments arose from RNase L cleavage, we analyzed transcriptome sequences immediately upstream of the 5’ mapped end of these mRNA fragment reads. In this case, we defined mRNA fragments as those shorter than one third of the length of the full-length annotated reference transcript that mapped to the 3’ end. It has been reported that the RNase L cleavage motif is UN^N ^21^ with preference for UU^N and UA^N based on reconstituted assays ^37^. Therefore, we expected enrichment of these motifs in our analysis. We focused on the region 10-15 nt upstream from the 5’ end of the reads due to the inability of the nanopores to sequence to the very 5’ end of mRNAs ^38^. In particular, we found that positions 11-12 nt upstream from the 5’ end of the read exhibited the highest information content for nucleotide preference as determined by bit depth (Figure 1D and S1G). Focusing on these positions, we found an enrichment of U and A nucleotides in 2-5A treated, but not untreated, samples. This preference is consistent with the view that mRNA fragment reads derive from mRNAs that are cleaved by RNase L. We also found that the proportional share of reads with U at position -12 increased in RNase L activated samples as compared to controls (Figure 1E and S1H), further supporting the idea that the detected mRNA fragments are the result of RNase L activation. Notably, we observed that UU, UA and UG motifs increased, while UC decreased in relative proportion, in agreement with previously described cleavage preferences in cell lysate ^21^. Interestingly, RNase L cleavage motifs in both WT and *XRN1* KO conditions were similarly detectable and showed comparable proportions within fragments (Figure 1 and S1). This suggests that XRN1 does not discriminate based on the origin of the fragments (RNase L cleaved vs those arising from other decay processes). These data indicate that the 3’ fragments left after mRNA cleavage by RNase L are readily detectable and abundant in the cell in comparison to full length transcripts.

### mRNA fragments generated by RNase L are bound by ribosomes

Based on evidence from ribosome profiling data, we previously proposed that ribosomes may initiate translation on mRNA fragments created by RNase L ^12,19^. To further test this model, we combined the above direct “long-read” nanopore sequencing method with sucrose gradient sedimentation to determine whether RNase L cleaved mRNA fragments were translated. Using this “polysome profiling” approach, fragments that are translated (or bound by stalled ribosomes) would be expected to migrate with ribosome-bound fractions in the gradient.

Since we detected high levels of RNase L cleaved mRNA fragments in *XRN1* KO cells, we used this cell line for these experiments to maximize the odds of detecting translation. *XRN1* KO cells were transfected with 2-5A (+2-5A, RNase L activator) or treated by the transfection reagent (−2-5A control) (Figure 2A). Then, we collected cell lysates and performed polysome profiling. We loaded lysate onto a 10-50% sucrose gradient that allowed separation of mRNAs loaded with ribosomes from the untranslated mRNAs by ultracentrifugation. Based on peaks in the UV absorbance reading, we collected fractions corresponding to monosomes, a single 80S ribosome, and polysomes (mRNAs bound to 2 or more ribosomes). The 80S fraction can include both “vacant” 80S ribosomes that lack mRNA and mRNAs bound to only one ribosome. In the control (−2-5A) lysates, we observed the expected distribution of ribosomes for unperturbed cells along the sucrose gradient with many ribosomes in both the “light” and “heavy” polysome fractions (Figure 2B). In contrast, in 2-5A treated samples, most ribosomes were found in the monosome (80S) fraction while the polysome fractions were drastically reduced with the heaviest polysomes eliminated entirely (Figure 2B) as compared to control cells. This loss of polysome peaks has been noted before ^11^ and likely occurs because mRNA levels in the cell are drastically reduced due to degradation, estimated at a level of 90% in previous reports where spike-in controls were utilized for normalization ^10,11^. Our observation that the most pronounced polysome peak in 2-5A treated samples is the first one, perhaps representing disomes (Figure 2B), suggests that fragments in the polysome pool are bound to a small number of ribosomes. We collected the small amount of total RNA present in the ribosome-bound fractions in both 2-5A treated and untreated cells and sequenced it by direct RNA nanopore sequencing as above (Figure 2A). We found that that both the monosome and polysome fractions contain 3’ mRNA fragments at levels similar to the total RNA control and far greater than respective −2-5A controls (Figure 2C and S2A-C). We also found similar RNase L cleavage motif patterns just upstream of the 5’ end of the fragments in the monosome and polysome fractions as in total RNA (Figure 2D-E, compare it to 1D-E). While we cannot precisely determine what proportion of the total population of 3’ fragments that are generated by RNase L get translated, the data suggest that at least some of these fragments can be loaded with one or a small number of ribosomes (or were perhaps already loaded prior to RNA cleavage) and are either engaged in active translation or stalled.

**Figure 2.**
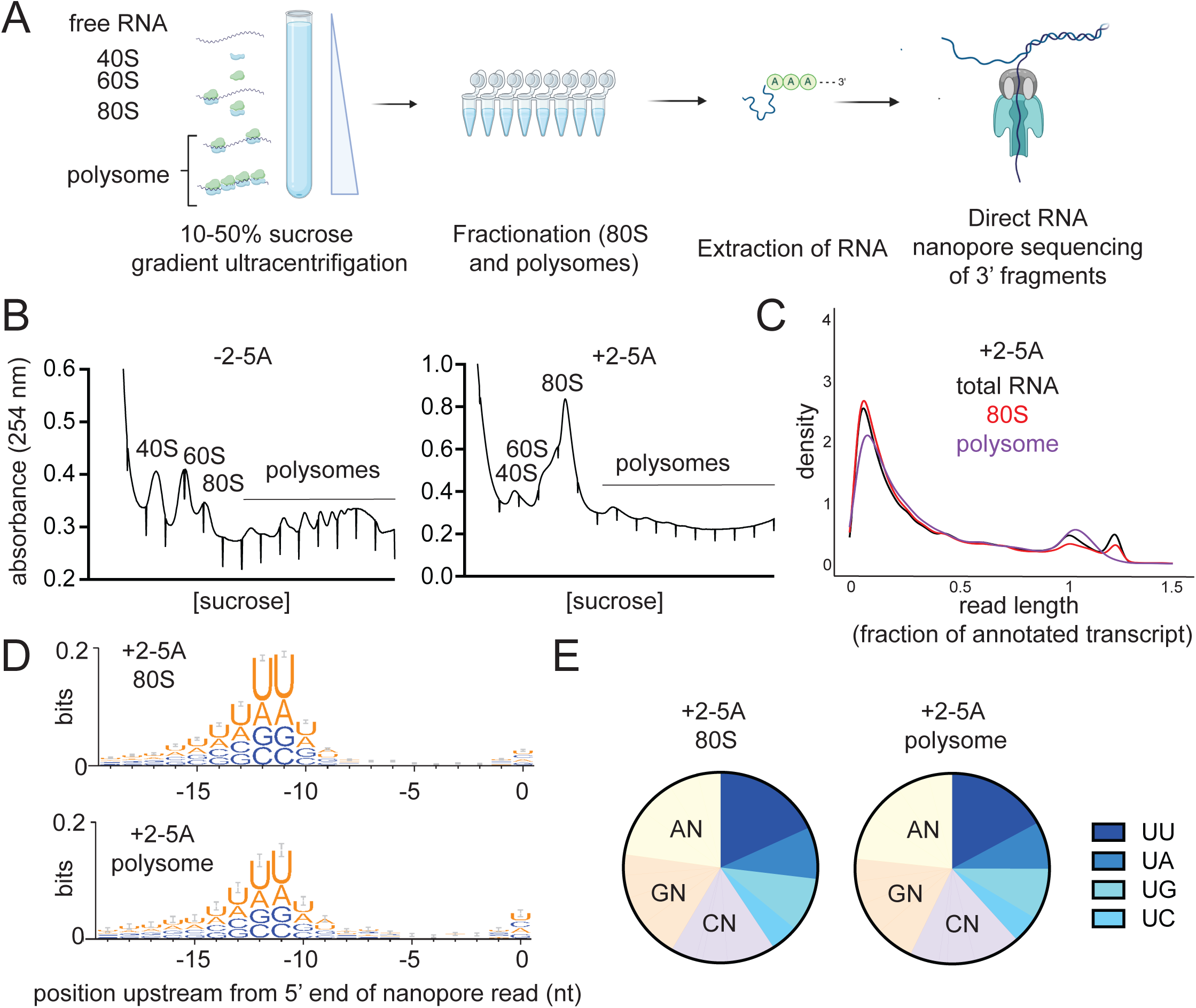
3’ fragments generated by RNase L are translated. **A** Schematic representation of polysome profiling experiments coupled with direct RNA nanopore sequencing. *XRN1* KO A549 cells are transfected with 5 µM of 2-5A for 4.5 hours. Then lysates were subjected to sucrose gradient centrifugation. 80S and polysome fractions were collected and RNA was extracted. Then, RNase L 3’ cleavage products and uncleaved mRNA that contained poly(A) were sequenced with direct RNA nanopore sequencing. **B** Polysome profiles of 2-5A treated (+2-5A, right panel) and untreated (−2-5A, left panel) cell lysates. RNA content was monitored by absorbance at 254 nm. Note that underlined area shows fractions retained for polysome analysis. **C** Normalized length distribution of nanopore direct sequencing reads to their respective annotated reference transcript in total RNA as compared to monosomes and polysomes in 2-5A treated *XRN1* KO cells shows fragmentation in all cases. **D** Weblogo analysis shows enrichment of U bases in transcriptome positions just upstream of where nanopore sequencing stopped in the 80S and polysome fractions of sucrose gradient sedimentation experiments in 2-5A treated *XRN1* KO cells. **E** Proportion of each di-nucleotide motifs near the 5’ end of 3’ fragments in 80S and polysome fractions of 2-5A treated *XRN1* KO cells. The number of total fragments were: 181,347 (80S) and 27,648 (polysome). In both D and E, 3’ fragments were defined as reads that were shorter than a third of their respective annotated transcript that mapped to the 3’ end.

### Deletion of the exonuclease XRN1 leads to increased mRNA fragment translation

As noted above, in our previous study we observed that the relative proportion of ribosome footprints increased in untranslated regions of mRNAs, including the 5’ and 3’ UTR regions as well as out-of-frame parts of coding sequences, as compared to the translated regions when RNase L was activated ^19^. This phenomenon was termed “altORF translation” because the footprints corresponded to distinct (alternate) open reading frames (ORFs) in these noncoding regions. Our interpretation of these data was that ribosomes could initiate on 3’ mRNA cleavage products and translate an open reading frame within the fragments^12,19^. Because the 3’ fragments are not capped (data not shown), ribosomes would have to initiate via a non-canonical process that would become more favorable due to mass action (high abundance of free ribosomes) in the cells under these conditions where cells lose the vast majority of their mRNA. Since we observed higher amounts of RNase L generated 3’ fragments in *XRN1* KO cells than in WT cells (Figure 1 and S1), it is expected that the KO cells should also exhibit a higher proportion of translation in altORFs than WT cells.

To test this hypothesis, we transfected WT and *XRN1* KO cells with 2-5A and performed ribosome profiling, a high-throughput ribosome footprinting approach that utilizes short-read Illumina sequencing (Figure 3A, see materials and methods). The RNase L activity levels were confirmed to be comparable by rRNA cleavage assay (Figure S3A). The most readily detected signature of altORF translation is a relative increase in ribosome footprints in the 3’UTR because this noncoding region of the mRNA is rarely translated by ribosomes under normal conditions ^19^. We used the ratio of 3’UTR to coding sequence (CDS) ribosome profiling reads to quantify the difference between control (−2-5A) and RNase L activated cells (+2-5A) and therefore assess the relative increase in altORF translation. We found that 2-5A treatment of *XRN1* KO cells led to a marked increase in the 3’UTR:CDS ratio as compared to WT cells (Figure 3B). This is consistent with a model where additional 3’ mRNA fragments in *XRN1* KO cells results in an increased proportion of ribosomes engaged in altORF translation. Additional analysis of averaged ribosome footprint levels (“metagene” analysis) further supported this interpretation. When we averaged the data in this way for all genes by aligning them by their stop codons (Figure 3C), we noted a higher level in the 3’UTR that is consistent with the above ratio-based analysis. We then performed metagene analysis on altORFs by aligning them by their start codons. This analysis provided strong evidence of translation, as exhibited by the 3-nt periodicity along the altORFs in 2-5A treated *XRN1* KO cells as compared to similarly treated WT cells (Figure 3D). RNase L activation also increases the proportion of ribosome footprints in 5’UTRs ^19^. We speculated that deletion of *XRN1* would not affect the share of these footprints since endonuclease-cleaved 5’ mRNA fragments are mostly degraded 3’ to 5’ by the exosome. In agreement with this prediction, we found that ribosome footprints in the 5’UTR were not elevated in *XRN1* KO cells as compared to WT during RNase L activation (Figure S3 B-D).

**Figure 3.**
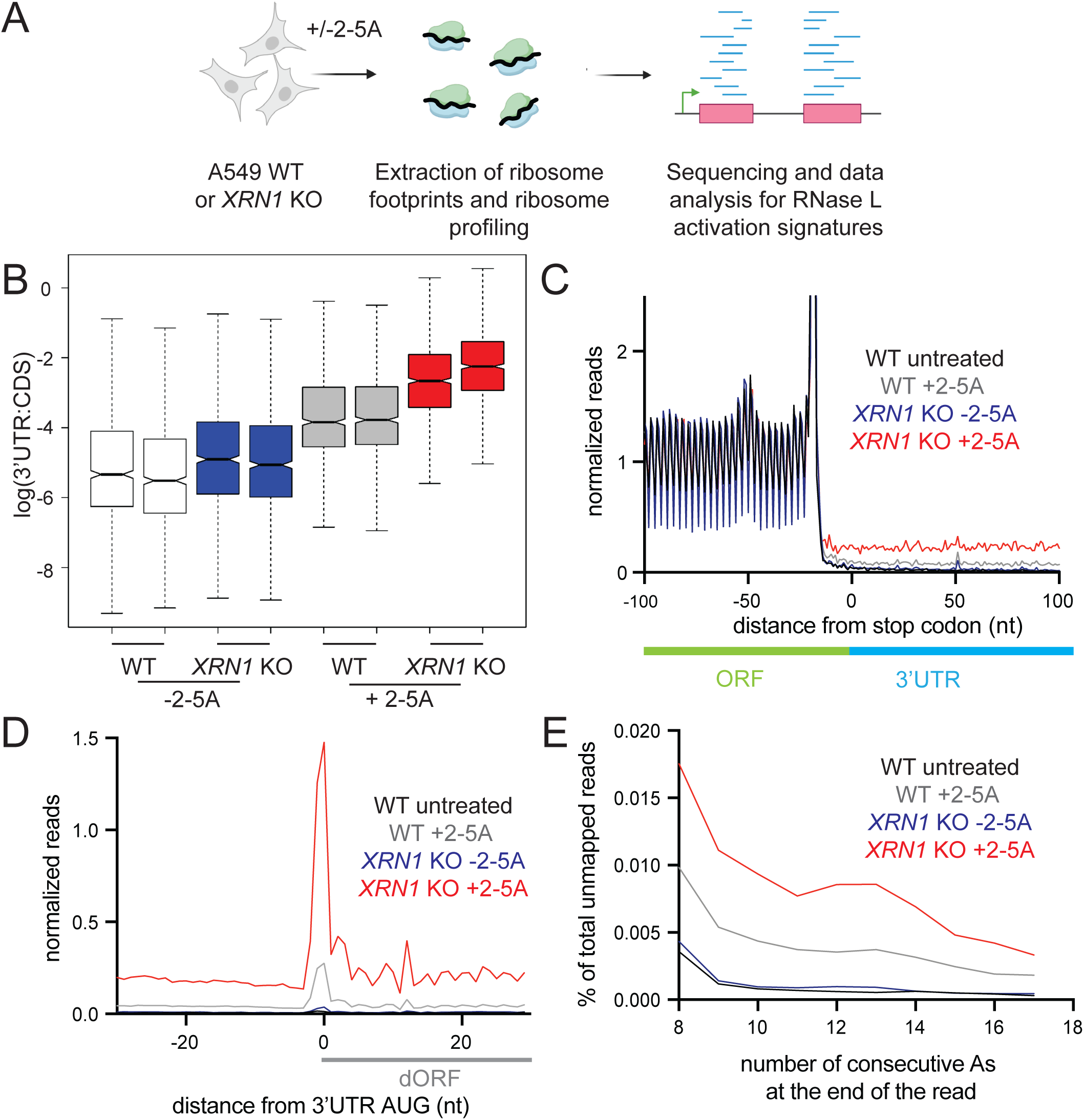
XRN1 KO cells exhibit stronger signatures of altORF translation compared to WT due to RNase L activation. **A** Schematic representation of ribosome profiling experiments. WT and *XRN1* KO A549 cells were treated with 5 µM 2-5A for 4.5 hours. Then lysates were used for ribosome profiling of 25-34 nt footprints. Data were analyzed for altORF translation signatures. **B** 3’UTR:CDS ratios increase in treated (+2-5A) and control (−2-5A) *XRN1* KO as compared to WT cells. **C** Normalized average ribosome footprint occupancy (metagene plot) around the stop codon of main ORFs (CDSs) reveals increased relative ribosome footprint levels in the 3’ UTRs when RNase L is activated vs the respective control (gray). *XRN1* KO cells further exhibit increase in 3’UTR footprints as compared to WT (red). Ribosome footprints are plotted by 5’ assignment without any shift. **D** Normalized average ribosome footprint occupancy around the start codon of downstream ORFs in the 3’UTR reveal 3-nt periodicity and increased translation during RNase L activation in *XRN1* KO cells as compared to WT cells. Ribosome footprints are plotted by 5’ assignment shifted by 12 nt (∼P site). **E** Percentage of ribosome profiling footprints in samples prior to mapping that contained consecutive poly(A) sequences (8-17 nt) at their 3’ end in WT and *XRN1* KO cells. These represent ribosomes that run into the poly(A) tail. The effect increases when XRN1 is absent.

An additional expectation of translation in 3’ UTRs is that, in some cases, ribosomes may not encounter a stop codon after initiating translation and would therefore translate the poly(A) tail. Translation of poly(A) is known to slow down or stall ribosomes engaged in elongation ^39–42^. In yeast, when conditions favor accumulation of ribosomes in poly(A) tails ^43^, ribosome footprints were found to protect a short stretch of poly(A) (8-15 nt) at the 3’ end of reads that did not map to the transcriptome since poly(A) tails are not encoded. To assess whether this occurred following RNase L activation, we quantified the percentage of reads that contain stretches of consecutive As at the 3’ end of ribosome profiling reads that did not map to the transcriptome (see Methods). We found the abundance of these reads was increased by RNase L activation as compared to control cells and further elevated in 2-5A treated *XRN1* KO cells (Figure 3E and S3E). These data are consistent with a model where RNase L activation leads to translation of the most distal 3’ mRNA fragments and results in ribosomes stalling in poly(A) tails of mRNAs.

### Ribosomes stall at RNase L cleavage sites

RNase L is known to activate ZAKα, a MAP3K that recognizes collided ribosomes ^12,13^. Based on this finding, we proposed that ribosomes stall and potentially collide at the 3’ ends of 5’ mRNA fragments that are created by RNase L. Prior work in yeast showed that such stalled ribosomes could be detected at cleavage sites that are generated by another endonuclease, Ire1 ^17^. Such footprints are distinct from ribosomes engaged in normal elongation because they protect a shorter footprint (∼16 nt vs ∼28 nt). These footprints are shorter since the cleaved mRNA 3’ end becomes positioned in the decoding center of the ribosome and no mRNA extends further into the mRNA entry channel (Figure 4A). Here, we used this short ribosome profiling method to detect stalled ribosomes at the 3’ ends created by RNase L cleavage. We treated A549 WT cells with 2-5A and performed modified ribosome profiling, where we selected for a wider range of footprints (15-34 nt), to capture short (15-18 nt, also termed “16-mer”) and full-length (25-34 nt, also called “28-mer”) ribosome footprints in the same sample (Figure 4A, see methods for details).

**Figure 4.**
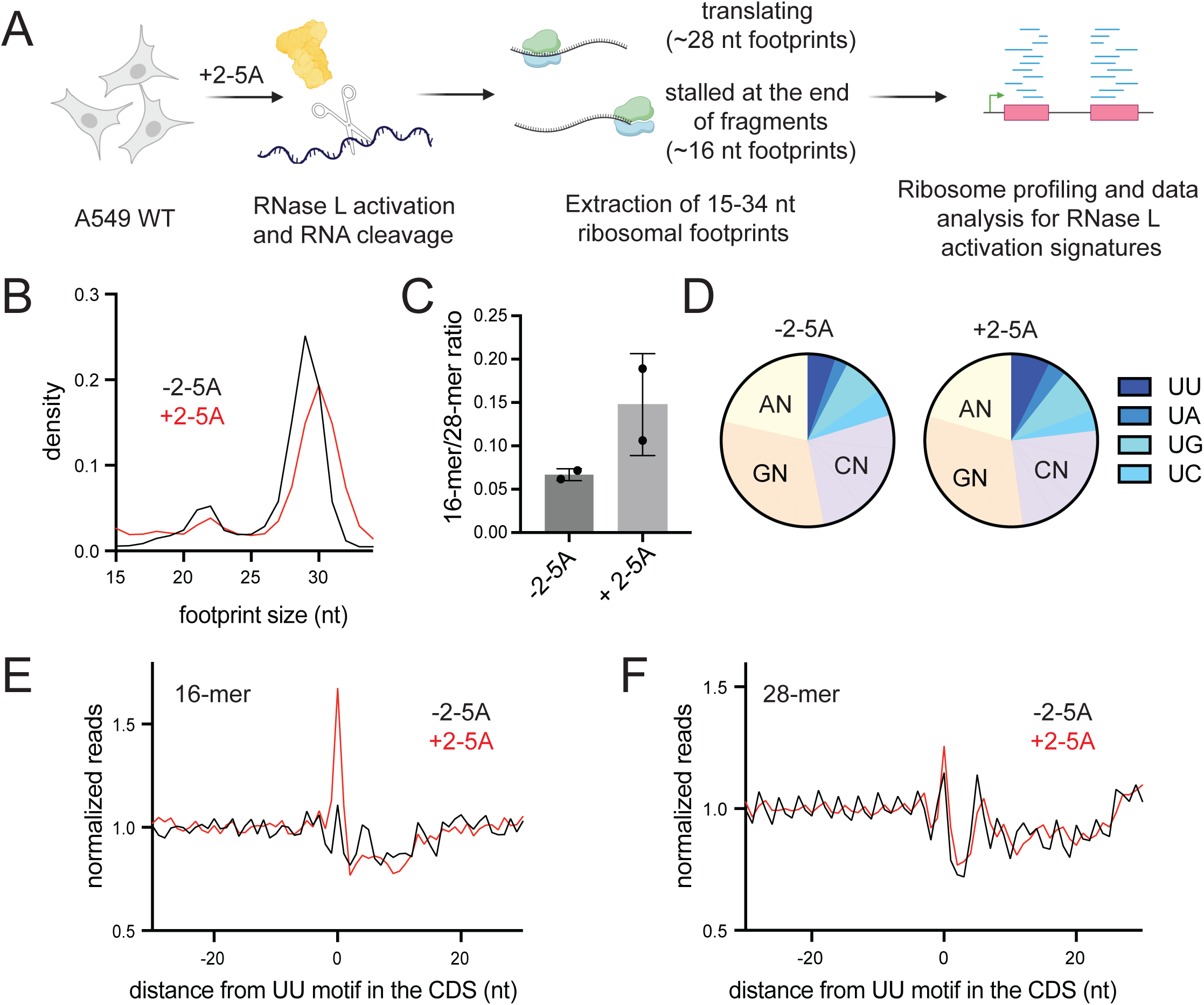
Short profiling footprints are increased at RNase L cleavage sites. **A** Schematic representation of ribosome profiling experiments. RNase L was activated by transfection of 2-5A for 4.5 hours and then cell lysates were subjected to ribosome profiling. We then performed size selection of 15-34 nt ribosome protected footprints to concurrently capture stalled ribosomes at the 3’ end of mRNA fragments (∼15-18 nt) and translating ribosomes (25-34 nt). Data then were analyzed to capture changes in length distribution of footprints and for RNase L signatures. **B** Normalized length distribution of transcriptome mapped ribosome profiling reads (15-34 nt). **C** Ratio of 16-mers (15-18 nt) to 28-mers (25-34 nt) in RNase L activated cells is increased. **D** 3’ end di-nucleotide motif distribution in 16-mer reads. Proportion of short footprints (15-18 nt) containing UU motif at their 3’ end is modestly increased in 2-5A treated cells. Number of 16-mer reads was 109,319 (-2-5A) and 333,784 (+2-5A). Position average plot of 3’ ends of 16-mers (**E**) and 28-mers (**F**) at RNase L cleavage sites (UU). 16-mers exhibit a peak at UU motifs when RNase L is active, consistent with ribosomes stalled on 3’ mRNA fragments. Ribosome footprints are plotted by 3’ assignment (no shift).

We found that RNase L activation shifted the size distribution of ribosome profiling footprints as the 16-mer footprints became proportionally more abundant in 2-5A treated cells than in control (Figure 4B and S4A). To quantify these changes, we also computed ratios of short and full-length footprints (“16-mer”/“28-mer” ratios). We found that the 16-mer/28-mer ratio increased upon RNase L activation, consistent with ribosomes stalling at the 3’ ends of cleaved 5’ mRNA fragments (Figure 4C). In addition, we note that a small 21-nt peak in the data decreased along with the 28-nt peak. This 21-nt peak is known to be a minor population of footprints and is created by ribosomes with an empty A site ^44^. The loss of this peak in tandem with the 28-nt peak lends support to our interpretation that the global population of 16-mer ribosome footprints increases in proportion to other species in the cell.

Since we observed an enrichment for UN^N cleavage motifs in transcripts just upstream of the sites where RNase L cleaved the RNA in our nanopore sequencing experiments (Figure 1), we examined the nature of the sequences just upstream of the 3’ end of the 16-mer footprints. We found that the 3’ dinucleotide sequence was somewhat more enriched in UN in 2-5A treated cells as compared to control (Figure 4D and S4B). To further study RNase L cleavage motifs in short footprints, we aligned the 3’ ends of 16-mer and 28-mer reads at UU motifs in the CDS to create an average plot (Figure 4E-F, S4C-D). We found a modest accumulation of 16-mers at UU motifs in 2-5A treated cells as compared to the control cells, suggesting enrichment of RNase L cleaved 3’ ends in short footprints. This further supports our model that ribosome stalling occurs at the RNase L cleaved ends of mRNA fragments. However, we note that these signatures of RNase L cleavage were less pronounced compared to those in nanopore sequencing data (Figure 1). This is expected because the footprints rely on a ribosome being stably associating with 3’ ends whereas the nanopore sequencing does not. Stable association of the ribosome on a 3’ end can be antagonized by ribosome rescue activity in the cell, particularly by PELO (see below).

As ZAKα is activated, we asked whether we could detect formation of queued ribosomes in the cell. Queued ribosomes behind 16-mers at Ire1 cleavage motifs were observed in yeast studies and supports a model where ribosomes can collide at the 3’ end of cleaved mRNA fragments^17^. However, in our data, we did not observe queueing of 28-mer footprints at RNase L motifs. This result suggests that either a single ribosome stalled at the end of the fragments is sufficient for activation of ZAKα or that the queued ribosomes are not particularly visible because they are tightly associated in disome structures and the RNase I used for ribosome footprinting cannot get between them. Another method that can reveal large changes in collided ribosome structures in cells is sucrose density gradient ultracentrifugation of lysates that are treated with an RNase to eliminate mRNA between ribosomes that are not immediate neighbors. We performed this experiment and observed a decrease in the disome peak relative to the 80S monosome peak in RNase L activated samples (Figure S4E). This result suggests that the absolute level of disomes decreases after 2-5A treatment. As a positive control, we treated cells with anisomycin, which is known to induce formation of disome structures that activate ZAKα^15^, and observed the expected increase the disome peak. However, it remains possible that a small population of (undetectable) disomes could form and activate this pathway. In particular, disomes are known to adopt different conformations ^45^ and this assay cannot distinguish between any change in their properties (and ability to be recognized by ZAKα) between 2-5A treated and control conditions. In addition, broad mRNA degradation by RNase L liberates ribosomes from mRNAs and therefore increases the number of empty 80S monosomes. This increase masks the true size of the mRNA-bound 80S monosome fraction, making it difficult to accurately compare the 80S peak to the disome peak in these cells. Therefore, the reduction in the generic disome peak here is inconclusive for determining the species that activates ZAKα.

### PELO rescues stalled ribosomes on mRNAs generated by RNase L activation

PELO is a rescue factor that can facilitate disassembly of stalled 80S ribosomes, with a preference for those at the 3’ end of endonuclease cleaved mRNAs ^17,43,46–48^. We therefore asked the question whether PELO has a role in recycling the ribosomes we observed to be stalled on the 3’ ends of 5’ cleavage fragments (Figure 5). We performed “knock-down” (KD) experiments against PELO combined with treatment of 2-5A in WT A549 cells. Then we performed ribosome profiling for 15-34 nt footprints (Figure 5A). The efficiency of PELO KD was monitored by western blotting and PELO protein levels were notably decreased (Figure 5B). The efficiency of RNase L activation was comparable across the samples (Figure S5A). We reasoned that reducing the amount of PELO would increase levels of ribosome stalling and collision, leading to increased activation of ZAKα the downstream MAP kinases JNK and p38. Thus, we monitored these kinases’ phosphorylation state using western blotting under RNase L activation. Indeed, we found that KD of PELO increased activation of ZAKα, JNK and p38 in 2-5A treated samples as compared to control siRNA treated cells (Figure 5B). Consistent with this, we found that short ribosome footprints also increased in PELO KD cells when RNase L was activate as compared to the control siRNA treated cells (Figure 5C-D, Figure S5B). Additionally, we observed an increase in the average level of 3’ ends of short footprints at UU motifs in average plots for short footprints in 2-5A treated PELO KD cells compared to WT (Figure 5E and S5C). Since the lack of a full knockdown and residual PELO in the cells could limit this trend, we additionally performed these experiments in HAP1 PELO KO cells where PELO was fully eliminated. We note that these cells are difficult to transfect with 2-5A and activation of 2-5A in rRNA cleavage assays is not prominent (Figure S5D). Regardless, we observed a stronger accumulation of 16-mer footprints at UU motifs in 2-5A treated PELO KO HAP1 cells than in A549 PELO KD cells (Figure 5F). We noted that 2-5A treatment did not increase average 16-mer footprints at UU motifs in WT HAP1 cells, presumably due to the low RNase L activation (Figure S5D). Our results suggest that PELO has a role in rescuing stalled ribosomes on mRNA fragments that are created by RNase L and may tune the innate immune response through the RSR.

**Figure 5.**
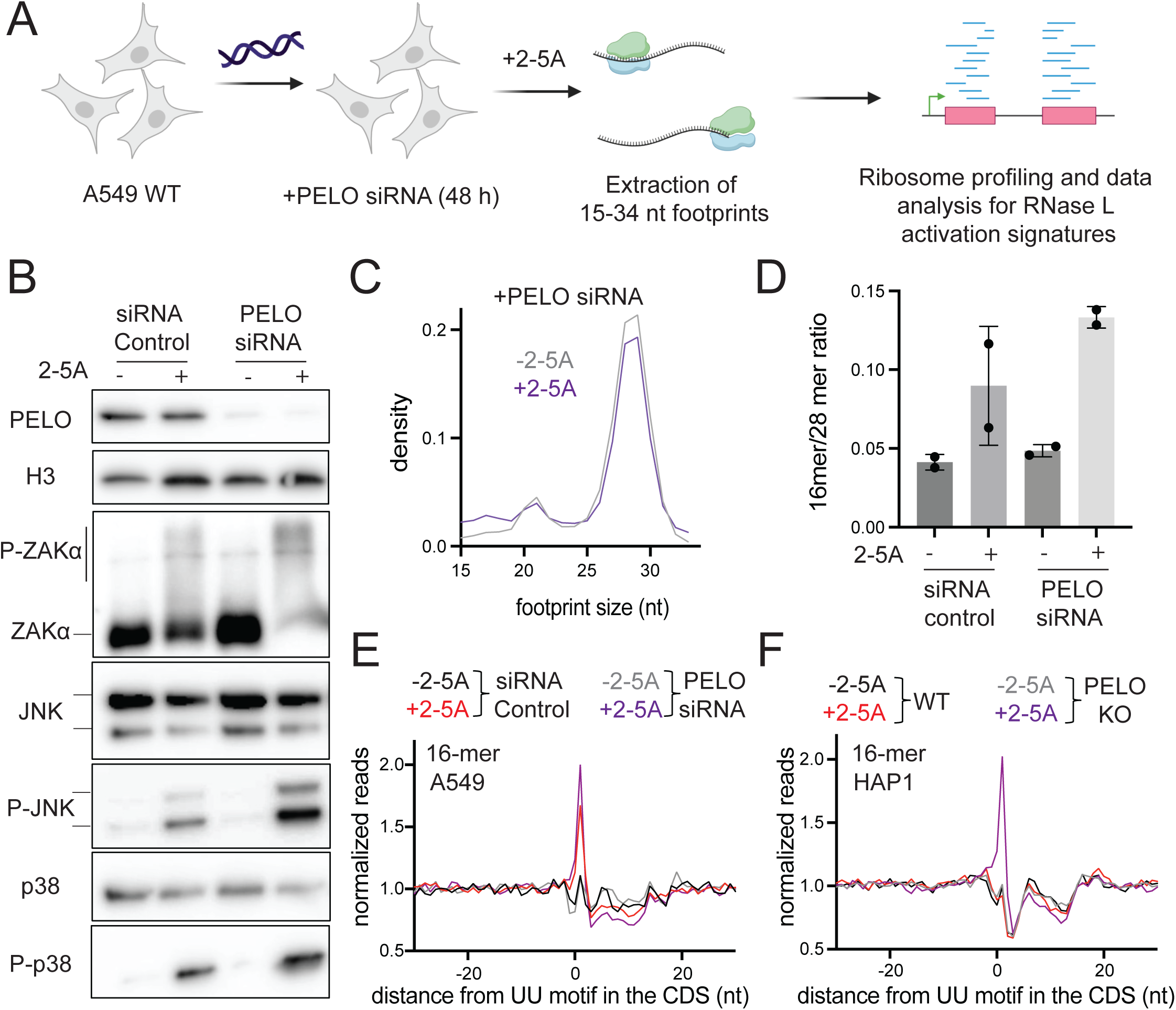
PELO rescues stalled ribosomes at the 3’ end of mRNA fragments during RNase L activation. **A** Schematic representation of experiments. A549 WT cells were treated with PELO siRNA or control siRNA for 48 hours followed by transfection with 2-5A or lipofectamine alone (−2-5A). Cells were collected and ribosome profiling for 15-34 nt footprints was carried out. **B** Western blots assay levels of protein during RNase L activation and PELO knockdown. Comparison of PELO abundance (upper panels) shows effectiveness of knock down. ZAKα phosphorylation, increased due to 2-5A treatment, was monitored by using western blotting on Phos-Tag gel. Shifts toward higher molecular weights indicate phosphorylation of ZAKα (middle panel). JNK and p38 phosphorylation (increased by 2-5A) were followed by western blotting with their respective phospho-antibody while also monitoring for total protein levels (lower panels). **C** Normalized length distribution of transcriptome mapped ribosome profiling reads (15-34 nt). **D** To analyze the effects of PELO knockdown, the ratio of 16-mers (15-18 nt) to 28-mers (25-34 nt) in PELO KD and RNase L activated cells was performed, demonstrating higher levels of 16-mers in PELO KD cells. Position average plot of 3’ ends of 16-mers near RNase L cleavage motif (UU) in PELO KD A549 cells (**E**) and PELO KO HAP1 cells (**F**).

## Discussion

A long-standing challenge is to understand how RNase L’s cleavage activity promotes death of infected cells and therefore immunity for the host organism. In this study, we established that the fragmented mRNA created by RNase L can be detected and that it is bound to ribosomes. The ribosomes translating these fragments can stall at 3’ ends of cleaved mRNAs. This stalling, and potentially collision with upstream ribosomes, activates ZAKα and the RSR, which promotes innate immunity by ultimately triggering cell death (Figure 6). Additionally, we found that the ribosome rescue factor PELO rescues stalled ribosomes at the 3’ ends of mRNA fragments and thus tunes the effects of RNase L’s downstream signaling (Figure 6).

**Figure 6.**
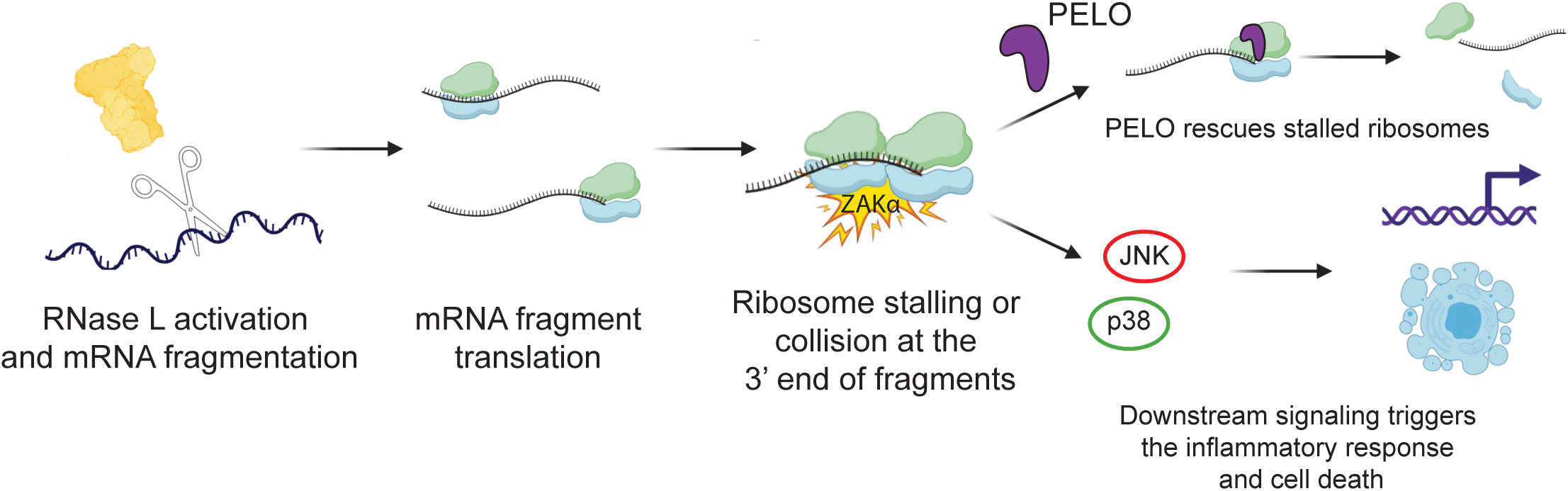
RNase L activation results in detectable amounts of mRNA fragments that are translated. Ribosomes stall at the end of the cleaved mRNA fragments and can be rescued by PELO (top pathway). They also induce activation of ZAKα and the downstream RSR via JNK and p38 and lead to the inflammatory response and cell death (bottom pathway).

Our use of direct nanopore sequencing showed that RNase L cleaved mRNA fragments increase in the cell and become even more abundant in cells lacking the 5’ to 3’ exonuclease, XRN1. Interestingly, the relative length of these species did not substantially change between WT and XRN1 KO cells,(Figure 1B-C; S1C-F), suggesting that once XRN1 engages with a fragmented mRNA, the degradation occurs rapidly. The observation that even under WT conditions (when XRN1 is present), a large number of RNase L cleaved RNA fragments persists suggests that the RNA decay machinery may be overwhelmed. It also possible that since RNase L leaves a 5’-OH on the 3’ cleavage fragment ^26,27^, there is some resistance to degradation because mRNA fragments with these ends are not ideal substrates for XRN1 ^30–32^. While these direct RNA nanopore sequencing experiments exclude studying mRNA fragments without a poly(A) tail (i.e. fragments made up of the 5’ end or intermediate segments of the mRNA), the development of alternative technologies to assess their abundance is needed in the future. In addition, it would be of interest to explore the stability of fragments created by other endonucleases, such as IRE1, SMG6, and others that have not yet been identified ^1,49,50^.

The stability of mRNA fragments that are generated by RNase L raises the question of whether they serve a function. It has been suggested that these fragments can form dsRNA and then bind RIG-I to induce the interferon production ^51^, but these effects have not universally been observed ^10,11,19^. Intriguingly, recent work showed that the loss of the 3’ to 5’ decay pathway via the exosome could enhance the transcription of genes that are sensitive to interferon, suggesting this function is tuned by the activity of this decay pathway ^28^. Another potential function for mRNA fragments is that they could be translated. We previously showed that altORFs within mRNAs are translated when RNase L is active ^19^ and one potential role of this translation is to produce short peptides that could act as non-self-epitopes presented on the cell surface and play a role in immunity. Translation of fragments also has the potential for a key signaling role. We and others recently showed that the ribosome collision sensor ZAKα^12,13^ is activated by RNase L and is the proximal cause for cell death ^14,15^.

Combination of our prior results, 1) that RNase L activity causes a proportional increase in translation of altORF sequences^19^ and 2) activation of the ZAKα ribosome collision sensor ^12,13^, indicate that RNase L mediates cellular responses via translation. However, the exact relationship between mRNA cleavage, translation, and ribosome stalling has been ambiguous. Our work here brings additional clarity to our understanding to the characteristics and outcomes of mRNA fragment translation. While activation of RNase L results in a sharp reduction in overall translation, as evidenced by the reduction in polysome peaks from a sucrose gradient, it is clear that the residual pool of mRNAs that is translated includes mRNA fragments, either by a single or small number of ribosomes.

How do ribosomes initiate translation on 3’ mRNA fragments that are cleaved by RNase L? RNase L cleavage leaves a 5’ OH, which is not favorable to initiation, unlike a 5’ cap. It is possible that some of these fragments are recapped by the cytoplasmic recapping machinery allowing efficient initiation on these fragments ^52^. However, we did not find evidence of for this based on CAGE-seq (cap analysis of gene expression) experiments (data not shown). Another possibility is that ribosomes can directly initiate on mRNA fragments ^19^ under conditions of RNase L activation. The loss of the vast majority of mRNA in the cell during RNase L activation will liberate most ribosomes from mRNA. This creates a ribosome abundance that could drive unfavorable initiation via mass action ^12,19^ that would load some mRNAs with one or a small number of ribosomes to an extent that we detected them here. This view is consistent with our previous observation that the proportion of altORFs that are translated during RNase activation increases as ribosomes presumably initiate on mRNA fragments and then translate nearby altORFs^19^. Consistent with this model, we showed here that this effect is augmented by the loss of XRN1 and the resultant increased abundance of these fragmented species in the cell.

While the presented nanopore sequencing results and previous ribosome profiling data strongly suggested that mRNA fragments can be translated by ribosomes, it is not clear how these ribosomes activate the ribosome collision sensor ZAKα. One possibility is that some altORFs within 3’ mRNA fragments lack a stop codon and instead lead into the poly(A) tail ^43^. Our data showed that the proportion of ribosomes that run into the poly(A) tail increased during RNase L activation and this effect was exacerbated in *XRN1* KO cells when compared to WT (Figure 3E). These ribosomes in the poly(A) tail may lead to ribosome collisions that activate ZAKα. Another possibility is that ribosomes translating the CDS or an altORF within a 5’ or internal mRNA fragment could encounter a 3’ cleaved end. In prior studies, ribosomes that encountered the cleavage sites of the endonuclease Ire1 protected a short (15-18 nt) footprint ^17^. Similarly, here we observed a global increase in the relative proportion of these footprints in cells where RNase L was active. While we did not directly observe an increase in queued ribosome footprints (Figure 4E) or an increase in RNase-insensitive disomes (Figure S4E) in these cells where RNase L is active, we cannot rule out the possibility that ribosome collisions capable of activating ZAKα occur at a low level or at an early timepoint.

We also asked the question of whether the ribosome stalling that is induced by RNase L could be modulated by the cell as a way to tune the downstream signaling to the RSR. We found that the reduction in the ribosome rescue factor PELO increased the level of stalling at RNase L’s UN^N cleavage sites and augmented the RSR. Given that SKIV2L is responsible for degradation of 5’ fragments, its elimination could also enhance stalling and the RSR. In this way, the ribosome rescue and decay machinery could be potentially tuned to regulate the innate immune response. Thus, developing inhibitors and or activators that target PELO could be considered for therapeutics. Interestingly, PELO was reported to directly associate with active RNase L to target exogenous RNA in the cell ^53^. How this interaction affects the efficiency or locus of RNase L cleavage activity remains an open question.

Our work establishes a model for how endonucleolytic cleavage by RNase L alters translation to evoke changes in key cellular pathways, such as cell death via the RSR. Since many endonucleases are involved in maintaining cell homeostasis and are important for responding to stress, it is possible that some of these findings apply to other endonucleases. It is also conceivable that endonucleases could be induced or added to cells for use as therapies to treat diseases such as cancer ^54^. In addition, viral endonucleases, such as coronavirus endonucleases^55^, are capable of broadly degrading host mRNAs. This suggests that the outcome of widespread endonucleolytic cleavage may not always be beneficial to the host. Addressing whether the activity of other endonucleases activate ZAKα or have other impacts on translation is a key question for the future with important implications for human health.

## Acknowledgements

We are thankful for the insightful discussions with Drs. Bret Hassel, Alan Hinnebusch, Jon Lorsch, and Tom Dever. We thank Dr. Bret Hassel and Dr. Emmanouil Maragkakis for feedback on the manuscript. We thank Dr. Sezen Meydan and Dr. Kyra Kerkhofs for helpful advice on short footprint ribosome profiling normalization and mapping. A549 cells were a kind gift from Dr. Bernie Moss. We also are thankful for assistance from the Genomics Core at the National Institute of Diabetes and Digestive and Kidney Diseases (NIDDK) and DNA Sequencing Core at the National Heart, Lung, and Blood Institute (NHLBI) for providing sequencing services for ribosome profiling. Additionally, we are grateful for the NHLBI core for allowing access to a GridION nanopore sequencer. We thank Dr. John Hanover for use of a FPLC. A.K. is grateful to have been chosen for a MOSAIC K99/R00 award (K99 GM143484) and for associated mentoring support from the American Society of Cell Biology partnership activities.

## Data availability

All raw sequencing (fastq) files are available at NCBI SRA in record PRJNA1269731 (ribosome profiling and direct nanopore RNA-seq). It can be accessed with the following link for reviewer access: https://dataview.ncbi.nlm.nih.gov/object/PRJNA1269731?reviewer=nll8rl17rob76depsbj01jnkd1 Custom code used for ribosome profiling and Nanopore sequencing in this study is available on Github: https://github.com/guydoshlab

## Author contributions

A.K. designed and performed experiments, developed software, analyzed the data and wrote the paper. G.D.J. performed HAP1 ribosome experiments related to Figure 5F and purified protein. N.R.G. designed experiments, developed software, and wrote the paper.

---

**Table 1.**
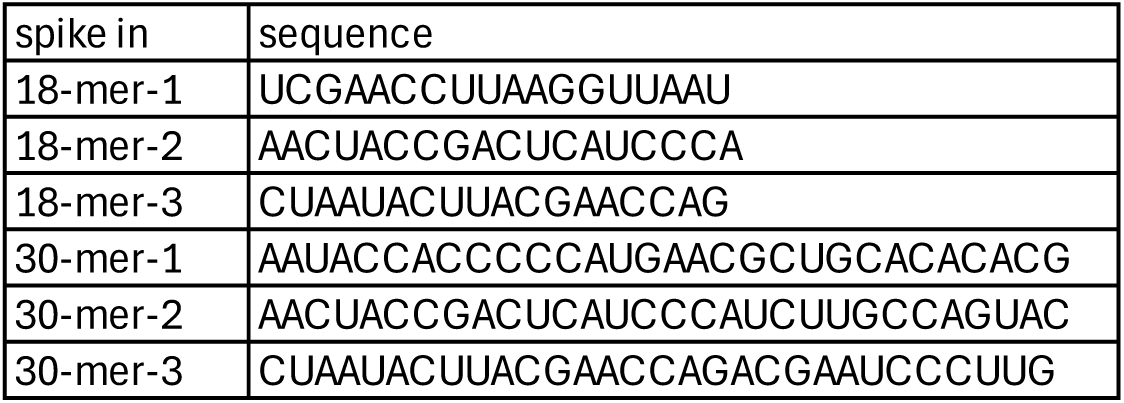
Oligonucleotide sequences utilized for spike-in normalization for 15-34 nt ribosome profiling experiments.

**Figure S1.**
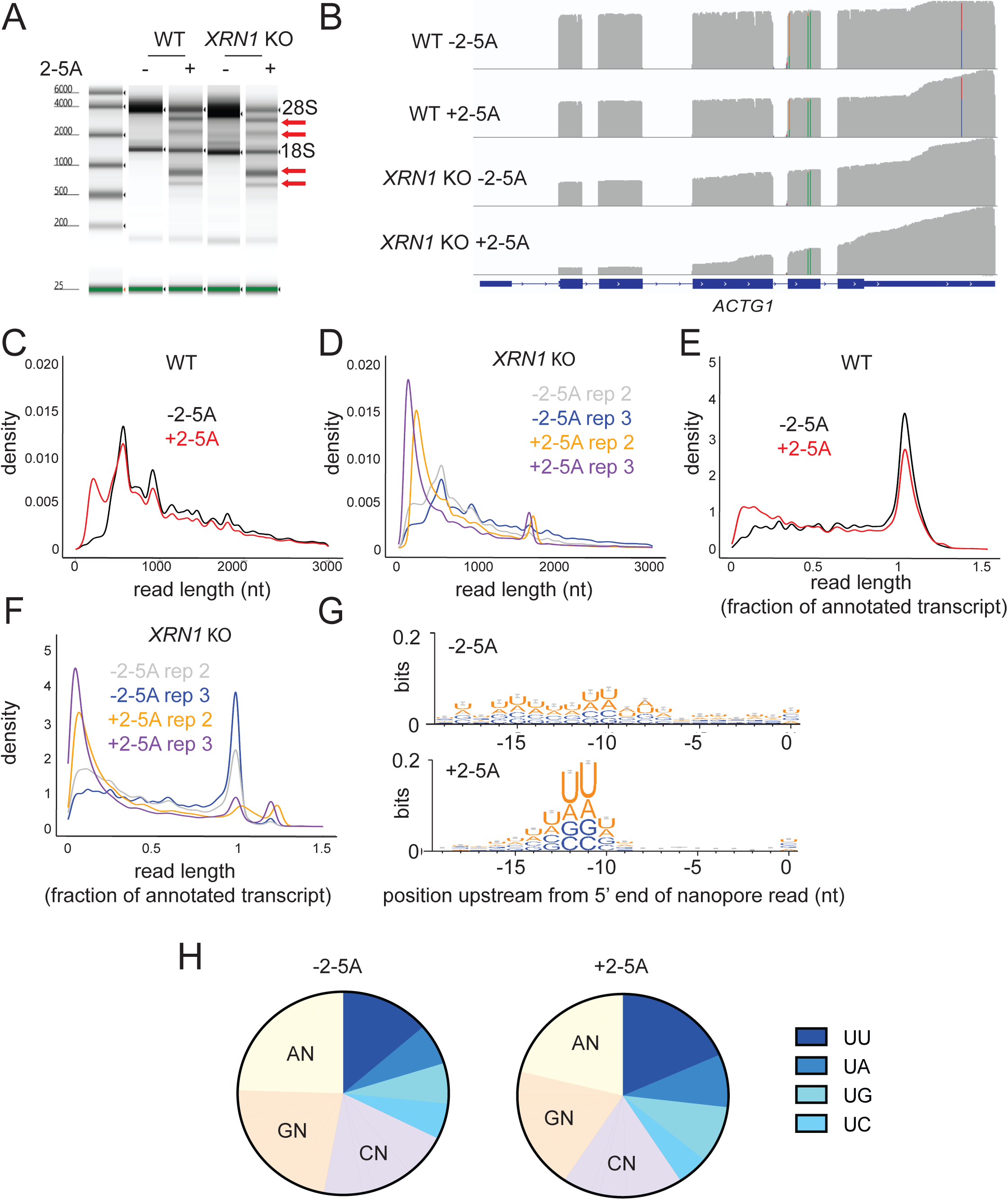
**A** rRNA cleavage assays showing RNase L activity in 2-5A treated but not in untreated cells. Red arrows indicate RNase L cleavage products. rRNA cleavage is comparable in WT and *XRN1* KO cells. Image output by Agilent TapeStation software with ladder shown on left. **B** Example gene model (*ACTG1*, oriented 5’ to 3’) showing a “pile up” view of nanopore raw (not normalized) reads. Image taken from IGV and colored bars indicate mismatches. **C** Length distribution of nanopore direct sequencing reads in 2-5A treated (+2-5A) and untreated (−2-5A) samples in WT cells. **D** Length distribution of nanopore direct sequencing reads in replicate 2-5A treated (+2-5A) and untreated (−2-5A) samples in *XRN1* KO cells. **E** Normalized length distribution of nanopore direct sequencing reads to their respective annotated reference transcript in 2-5A treated (+2-5A) and untreated (−2-5A) samples in WT cells. **F** Normalized length distribution of replicate nanopore direct sequencing reads to their respective annotated reference transcript in 2-5A treated (+2-5A) and untreated (−2-5A) samples in *XRN1* KO cells. **G** Weblogo analysis shows enrichment of U bases in transcriptome positions just upstream of where nanopore sequencing stopped in ±2-5A treated WT cells. **H** Di-nucleotide motif distribution near the 5’ end of 3’ fragments in ±2-5A treated WT cells (−11-−12 positions as shown in G to account for RNA left in pore after sequencing stops). Total number of fragments in each sample were: 110,068 (-2-5A), 619,964 (+2-5A). In both G and H, 3’ fragments were defined as reads that were shorter than a third of their respective annotated transcript.

**Figure S2.**
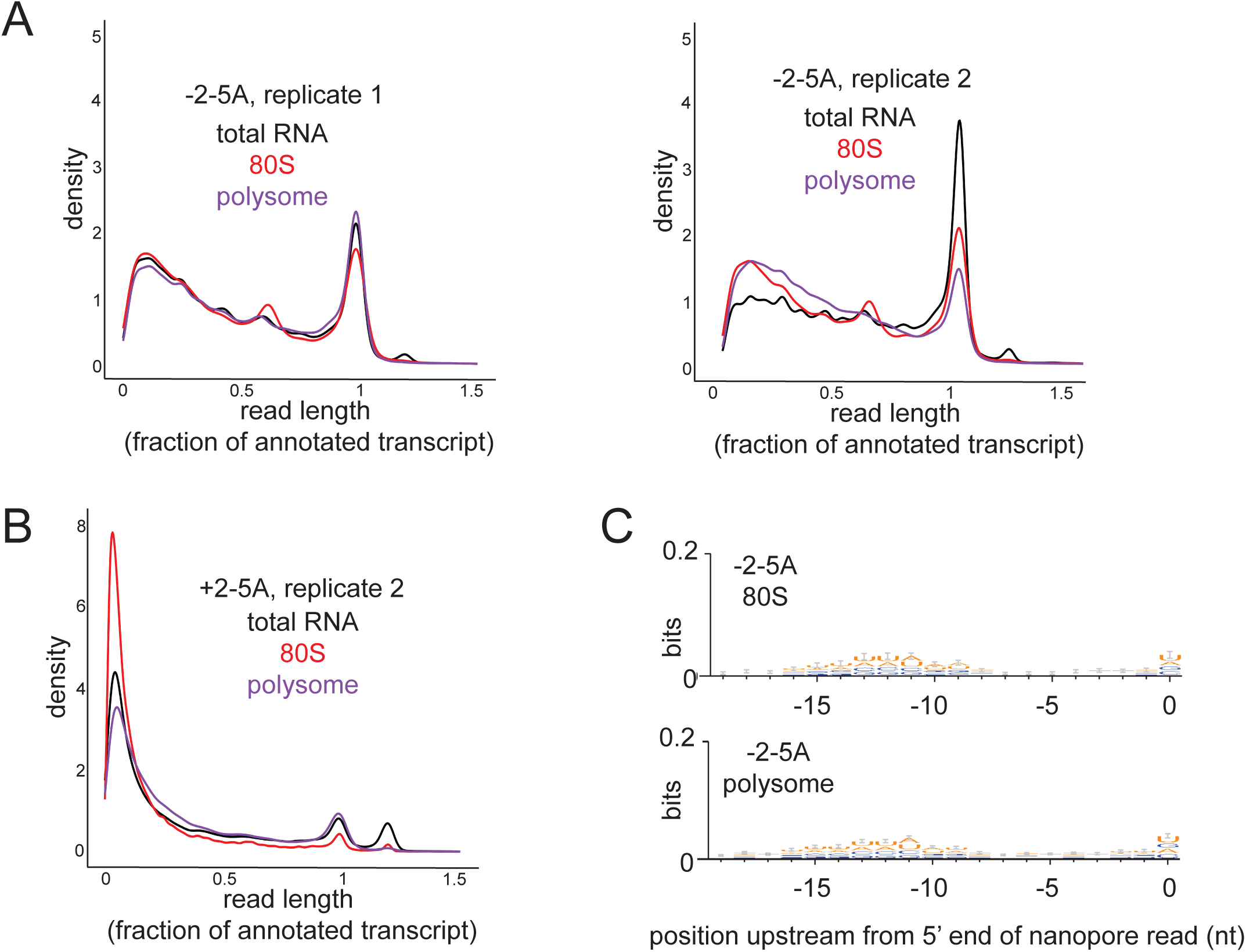
**A** Normalized length distribution of nanopore direct sequencing reads to the annotated reference transcript in total RNA, monosomes and polysomes in untreated *XRN1* KO cells (replicate 1 and 2). **B** Replicate experiments for Figure 2C. **C** Weblogo analysis shows enrichment of U bases in transcriptome positions just upstream of where nanopore sequencing stopped in the 80S and polysome fractions of sucrose gradient sedimentation experiments in untreated *XRN1* KO cells.

**Figure S3.**
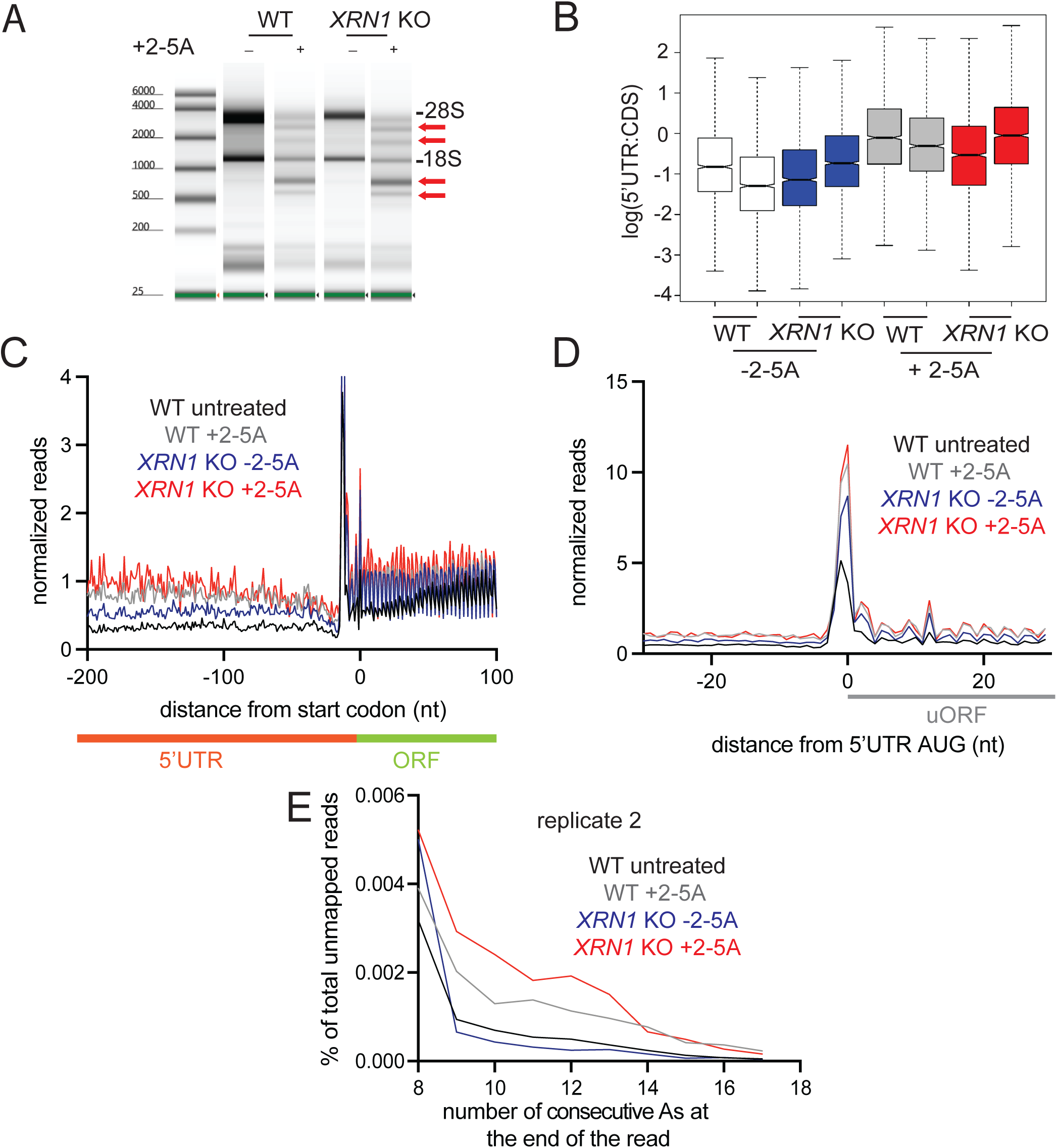
**A** rRNA cleavage assays in lysates used in ribosome profiling showing RNase L activity in 2-5A treated but not in untreated cells (replicate 1). Red arrows indicate RNase L cleavage products. rRNA cleavage is comparable in WT and *XRN1* KO cells. Image output by Agilent TapeStation software with ladder shown on left. **B** 5’UTR:CDS ratios do not increase in treated (+2-5A) and control (−2-5A) *XRN1* KO as compared to WT cells. **C** Normalized average ribosome footprint occupancy (metagene plot) around the start codon of main ORFs (CDSs) reveals increased relative ribosome footprint levels in 5’ UTRs when RNase L is active vs −2-5A control. XRN1 KO cells don’t exhibit increase in 5’UTR footprints as compared to WT (red). Ribosome footprints are plotted by 5’ assignment without any shift. **D** Normalized average ribosome footprint occupancy around the start codon of upstream ORFs in the 5’UTR reveal increased uORF translation during RNase L activation. This is not further increased in *XRN1* KO cells as compared to WT cells. Ribosome footprints are plotted by 5’ assignment shifted by 12 nt (∼P site). **E** Percentage of ribosome profiling footprints in samples prior to mapping that contained consecutive poly(A) sequences (8-17 nt) at their 3’ end in WT and *XRN1* KO cells (replicate 2). These represent ribosomes that run into the poly(A) tail. The effect increases when XRN1 is absent.

**Figure S4.**
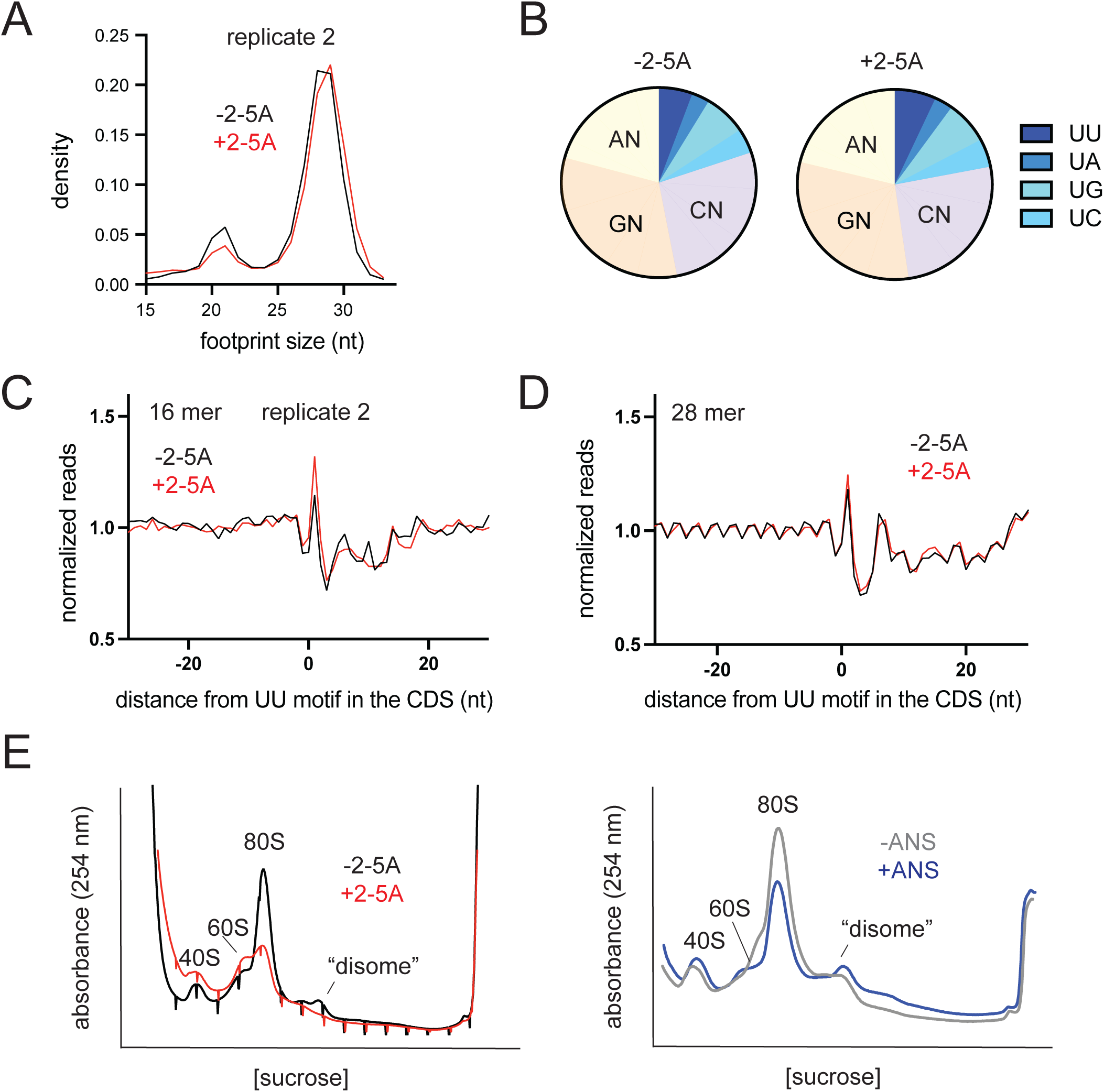
**A** Normalized length distribution of transcriptome mapped ribosome profiling reads (15-34 nt) for replicate experiments (replicate 2). **B** 3’ end di-nucleotide motif distribution in 16-mer reads. Proportion of short footprints (15-18 nt) containing UU motif at their 3’ end is modestly increased in 2-5A treated cells (replicate 2). **D** and **C** Position average plot of 3’ ends of 16-mers and 28-mers at RNase L cleavage sites (UU) for replicate experiments. Ribosome footprints are plotted by 3’ assignment. **E** RNase A treatment of the lysates combined with 10-35% sucrose gradient ultracentrifugation serves as an assay for ribosome collisions with ANS serving as a positive control. The level of RNA was monitored by absorbance at 254 nm during fractionation.

**Figure S5.**
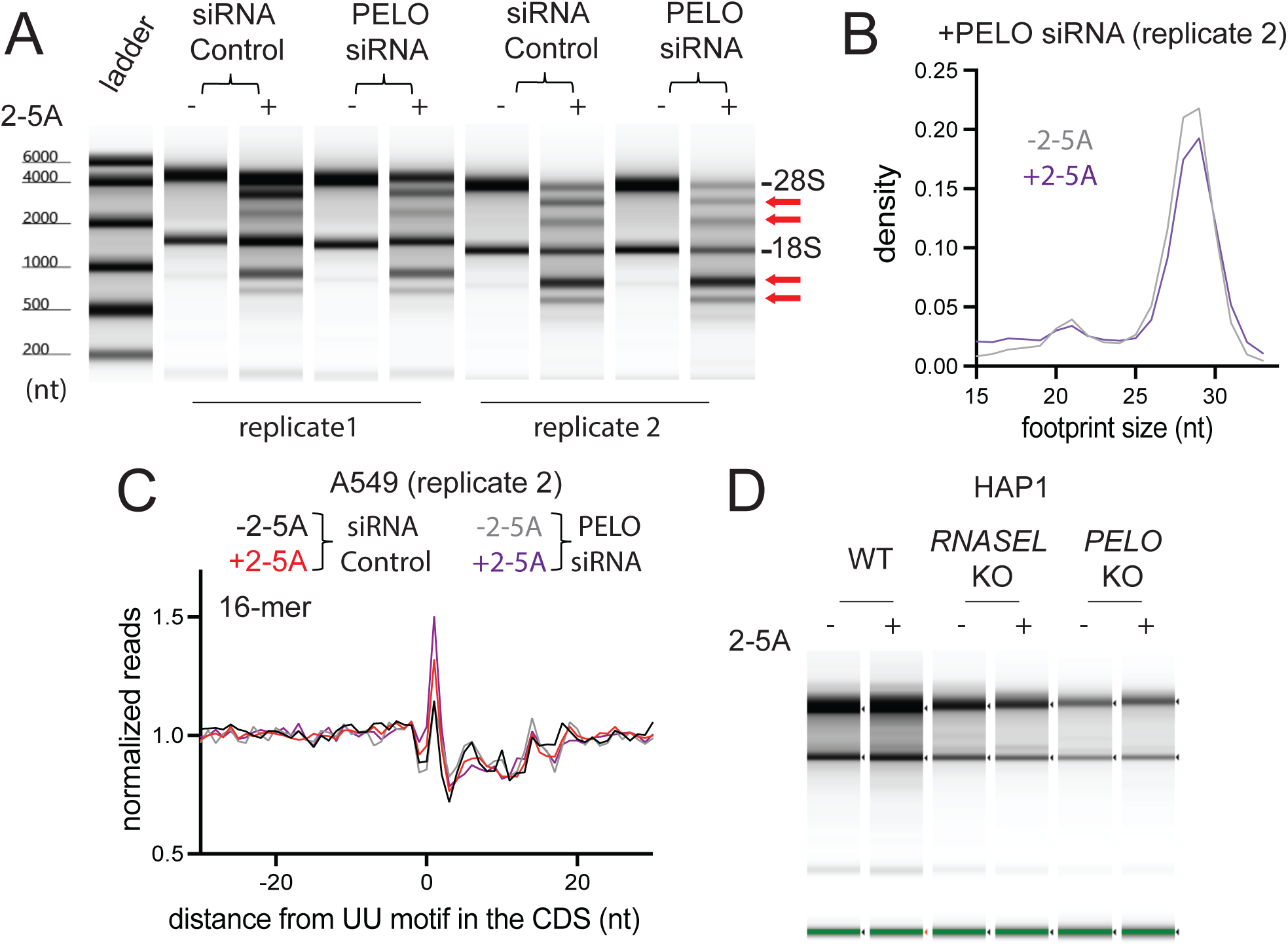
**A** rRNA cleavage assays from lysates used in the 15-34 nt ribosome profiling experiments where PELO was knocked down show RNase L activity in 2-5A treated but not in untreated cells. Red arrows indicate RNase L cleavage products. rRNA cleavage is comparable across samples. **B** Normalized length distribution of transcriptome mapped ribosome profiling reads (15-34 nt) in PELO siRNA treated cells (replicate 2). **C** Position average plot of 3’ ends of 16-mers near RNase L cleavage motif (UU) in PELO KD A549 cells (replicate 2). **D** rRNA cleavage assays in HAP1 WT and PELO KO cells shows little RNase L activity.

## Materials and Methods

### Cell culture

WT and *XRN1* KO A549 lung carcinoma cells were cultured in RPMI (Gibco Cat # 60870127) complemented with 10% Fetal Bovine Serum (Gibco, A56704). HAP1 WT and PELO KO cells were cultured in IMEM RPMI complemented with 10% Fetal Bovine Serum (Gibco, A56704). Cells were tested and negative for Mycoplasma contamination throughout the study. Mycoplasma testing was performed using eMyco Valid Mycoplasma PCR detection kit (BioLink) or ATCC Universal Mycoplasma Detection Kit. Cells were incubated at 37°C in the presence of 5% CO_2_. A549 cell lines used in this study were a kind gift of Dr. Bernie Moss (NIH) and generated as described in ^56^. HAP1 WT and PELO KO cell lines were purchased form Horizon.

### rRNA cleavage assay

Total RNA was extracted from ∼10^6^ cells or 30-50 µl ribosome profiling cell lysates using RNeasy kit (Qiagen) according to the manufacturer’s protocol. The amount of total RNA was computed by absorbance at 260 nm measured by NanoDrop spectrophotometer (Thermo Fischer Scientific) and then diluted to 50-200 ng/µl. Then RNA samples were run on a TapeStation 4150 with the Agilent RNA ScreenTape assay. Data visualization was done by Tape Station Analysis Software 4.1.1.

### Synthesis and purification of 2-5A

2-5A was synthesized by recombinant human OAS1 in an *in vitro* assay as described previously^12,19^. First, recombinant OAS1 (p42) containing an N-terminal His tag was expressed in BL21(DE3) *E.coli* as described before ^57^ or with an alternative protocol using autoinduction media ^58^. Cells were pelleted by centrifugation at 7000 g for 15 min at 4°C and lysed in B-per protein extraction reagent (4 ml/gram bacterial pellet) in the presence of cOmplete mini protease inhibitor cocktail (Roche) for 15 minutes at room temperature. Next, bacterial lysate was cleared by centrifugation at 34,000 g for 1 hour at 4°C. The supernatant was filtered (45 µm pore size, Millipore) and loaded onto a HisTrap (GE Healthcare) nickel column. The column was washed with wash buffer (20 mM Hepes pH 7.5, 300 mM NaCl, 10% (vol/vol) glycerol, 1 mM TCEP and 50 mM imidazole) and OAS1 was gradient eluted with 500 mM imidazole. Protein fractions were then evaluated by SDS-PAGE and Coomassie staining. Then OAS1 containing fractions were pooled, concentrated and buffer exchanged (Zeba Spin desalting Column, 7 K MWCO, Thermo Scientific) in storage buffer (20 mM Hepes (pH 7.5), 300 mM NaCl, 10% (vol/vol) glycerol and 1 mM TCEP). Final protein preparation was stored in storage buffer at -80°C. Concentration of OAS1 was determined by using NanoDrop Spectrophotometer (Thermo Fischer Scientific) (Mw=41.5 g/mol and ε= 65,485 M^-1^cm^-1^).

To produce 2-5A, 2 µM purified OAS1 was incubated with 1.25 OD_260_ poly I:C in the presence of 10 mM ATP, 20 mM Hepes (pH 7.5), 50 mM NaCl, 30 mM MgCl_2_, 10% (vol/vol) glycerol, 4 mM DTT at 30°C for 2 hours. To stop the reaction, samples were incubated at 85°C for 15 minutes. Then the samples were filtered (22 µm pore size Millipore filter) and the different 2-5A species were separated on a 16/10 MonoQ column as described before ^57^. The same 2-5A fractions from several runs were pooled and run again on a 16/10 MonoQ column to achieve higher concentrations. 2-5A concertation was estimated by NanoDrop Spectrophotometer at 259 nm. Then yielded 2-5A was aliquoted and stored at -80°C.

### Polysome profiling via sucrose gradient sedimentation

*XRN1* KO A549 cells were grown to ∼70% confluency in a T75 flask (∼8×10^6^ cells) and were transfected with 5 µM 2-5A or transfection reagent only (control) for 4.5 hours with Lipofectamine 3000 as outlined in the manufacturer’s protocol. After 4.5 hours of incubation at 37°C, cells were treated with media containing 50 µg/ml of cycloheximide for 10 minutes at 37°C. Cells were then washed twice with ice-cold DPBS containing 50 µg/ml of cycloheximide before the addition of 600 µl of lysate buffer (20 mM Tris (pH 8.0), 140 mM KCl, 5 mM MgCl_2_, 0.5 mM DTT, 0.03% Nonidet P-40, 100 mg/ml cycloheximide, 300 U/ml Superasein, and EDTA-free protease inhibitor cocktail tablet (Roche)). Cells were scraped from the bottom of the plates and were transferred to a microcentrifuge tube. Then cells were passed through a G-26 needle for 10 times and incubated on ice for 10 minutes. Lysates were cleared by centrifugation at 4°C, 8,000 g for 10 minutes and the supernatants were transferred to a new tube. Samples were flash frozen in liquid nitrogen and stored at −80°C until the day of sucrose gradient centrifugation. 10-50 % sucrose gradients were made fresh before the gradient ultracentrifugation. For this, 60% sucrose was prepared and filtered through a 0.22 µm filter. Then 10% and 50% sucrose gradient solutions were diluted with molecular grade water from this 60% sucrose and 10X gradient solution (200 mM Tris (pH 8.0), 1.5 M KCl, 50 mM MgCl_2_ and 5 mM DTT, to be diluted to 1X). To make the sucrose gradients, polypropylene centrifuge tubes (Beckman coulter, 14X89 mm, #331372) were first filled with 6 ml of 10% sucrose gradient buffer and ∼6 ml of 50% sucrose gradient solution was underlaid using a canula attached to a syringe. Gradients were made using a tilted tube rotation method (Biocomp) with a standard program for 10-50% gradients (1 min 48 sec, 81.5°, 17 rpm). The top 1 ml of the gradients was discarded, and cell lysates were carefully loaded dropwise onto the top. Then gradients loaded with samples were centrifuged in a swinging rotor (SW41Ti, Beckman Coulter) at 40,000 rpm for 2 hours at 4°C. Then the gradients were fractionated using a Brandel Density Gradient Fractionation System while the RNA content was monitored with at 254 nm and digitally recorded with a DataQ Instruments DI-155. Fractions were stored at -80°C or used immediately for RNA precipitation.

### RNase digestion and sucrose gradient

WT A549 cells were grown to ∼70-80% confluency in a T75 flask (∼8×10^6^ cells) and were transfected with 5 µM 2-5A or transfection reagent only (control) for 4.5 hours with Lipofectamine 3000 as outlined in the manufacturer’s protocol or treated with 1 mg/ml anisomycin diluted in 10 ml of media (Sigma, #A9789). This concentration of anisomycin is known to induce strong disome formation and thus acted as a positive control in our experiments. After 4.5 hours (2-5A treatment) or 15 minutes (anisomycin treatment) of incubation at 37°C, cells were washed with 10 ml room temperature DPBS twice, then lysed in 600 µl lysis buffer (20 mM Tris-HCl, pH=8.0, 150 mM NaCl, 15 mM MgCl_2_, 1% Triton-X, 1 mM DTT, 1X phosphatase inhibitor cocktail (Cell Signaling), 1X EDTA-free protease inhibitor cocktail (Roche)) while scraped off from the bottom of the plate. Lysates were incubated for 10 minutes on ice and then homogenized by pushing it through a G-26 needle before proceeding for centrifugation (8,000 g, 10 minutes at 4°C). Total RNA was quantified by Quant-IT RiboGreen kit (Invitrogen, R11490) using a mini-fluorimeter (Turner Biosystems, TBS-380). Lysates containing 20-100 µg of RNA were diluted to 800 µl with lysis buffer and 2.66 µl of 0.5 µg/µl RNase A (Thermo Fischer Scientific, #EN05310) was added. RNase A digestion was carried out for 15 minutes at 25°C, while shaking the tubes at 700 rpm. Lysates then were loaded onto 10-35% sucrose gradients, prepared similarly to above (see polysome profiling), except that the final concentration of the gradient buffer contained 20 mM Tris-HCl, pH=8.0, 150 mM NaCl, 50 mM MgCl_2_. Ultracentrifugation and fractionation steps were carried out similar to the procedure described for polysome profiling.

### Nanopore sequencing

#### Sample and RNA library preparation

WT and *XRN1* KO A549 cells were grown to ∼70% confluency in a T75 flask (∼8×10^6^) and were transfected with 5 µM 2-5A for 4.5 hours with Lipofectamine 3000 (Invitrogen) as outlined in the manufacturer’s protocol. We used Lipofectamine treated cells as controls (no 2-5A) in these experiments. Cells were collected by centrifugation and total RNA was extracted with Qiagen RNeasy kit. For each sample RNase L activation was monitored by rRNA cleavage assays (above). Total RNA for samples from polysome profiling experiments (Figure 2) were precipitated from appropriate fractions as obtained above (see polysome profiling). To each polysome profiling fraction (∼500 µl), we added 2 volumes of ethanol (1 ml), 40 µl of 1 M sodium acetate (pH=5.5) and 1 µl of Glycoblue (Invitrogen, AM9516) and precipitated RNA at -20°C overnight. The next day the precipitated RNA was collected by centrifugation at 4°C, 20,000 g for 30-60 minutes until a pellet was visible at the bottom or the side of the tube. Then the pellet was washed with 70% ethanol, dried for ∼10 minutes and resuspended in 10-15 µl of water. Total RNA was quantified by Quant-IT RiboGreen kit (Invitrogen, R11490) using a mini-fluorimeter (Turner Biosystems, TBS-380). Enrichment for polyadenylated mRNA fragments were carried out by Invitrogen Dynabeads mRNA purification kit (#61006). Then poly(A) enriched (150-300 ng) or total RNA (0.5-1 µg) was used for creating libraries for direct RNA nanopore sequencing. Library preparation was carried out with Nanopore direct RNA sequencing kit (SQKRNA004) according to the manufacturer’s protocol, including the reverse transcription step to facilitate RNA sequencing. We used 0.1-0.5 µl of control mRNA (labeled “RNA CS” in the kit) to ensure proper library preparation and to monitor potential problems during sequencing. Sequencing was performed on MinION flow cells (FLO-MIN004RA) using a GridION sequencer. Fastq files containing the long read sequences were generated by MinKNOW 6.2.14 software using default settings (min quality score= 9) for direct RNA sequencing.

#### Data processing and analysis

Fastq files were aligned to the human transcriptome (Ref-Seq Select+MANE, ncbiRefSeqSelect, as downloaded from UCSC on 14 April 2020) or the human genome (hg38, UCSC, only for Figure S1B) by minimap2. Duplicate entries (using SeqKit software rmdup function) were removed prior alignment. For transcriptome and genome alignment, we aligned the reads using a preset designed for long reads (-ax map-ont) without secondary alignments (--secondary=no).

For monitoring normalized length distribution of human transcriptome mapped reads and to extract nucleotide sequences for motif analysis we used custom python scripts (see Guydoshlab github page). To generate the weblogo and di-nucleotide motif distribution plots we used a length range of 50 bases and 10 kb and required the read to be <1/3 of total annotated length mapping at the 3’ end. Nanopore length density plots (Figures 1-2, S1-S2) were generated with the geom_density function and plotted by ggplot2 in R and R Studio (version 2024.12.1). To visualize nucleotide enrichment near the end of 5’ ends of 3’ fragments we used weblogo 3.7.12. Distribution of nucleotide motifs were plotted with Prism 10.2.2. Differences between length distributions in ±2-5A treated cells were assessed using Whitney-Mann-Wilcoxon test. Assigned p values were lower than 0.01 for all comparisons.

### siRNA treatment of A549 cells

WT A549 cells were grown to ∼70-80% confluency in a T75 flask (∼8×10^6^ cells) and transfected with PELO (IDT, hs.Ri.PELO.13.2) or control siRNA (IDT, #51-01-14-03) using Lipofectamine 3000 transfection reagent according to the manufacturer’s protocol. After 24 hours cells were trypsinized and plated to achieve ∼40% confluency. siRNA treated cells were incubated for another 24 hours (total treatment is for 48 hours) before transfection with 2-5A (5 µM) or transfection reagent only (Lipofectamine 3000, Invitrogen). Cells were then incubated for an additional 4.5 hours before their lysis for ribosome profiling experiments. Decrease in PELO protein levels was confirmed by western blotting.

### Ribosome profiling

#### Sample and cDNA library preparation

Ribosome profiling of “conventional” 25-34 nt footprints was carried out as previously reported^19^, except that rRNA depletion was carried out by siTools riboPool rRNA depletion kit designed for human ribosome profiling (Figure 3). For data in Figure 4, we selected a broader range of footprints (15-34 nt) during the size selection step using an additional 15 nt RNA marker ^17^. Additionally, we mixed 18 and 30 nt “spike-in” RNAs (3 different sequences each) with ribosome protected footprints during the size selection step to control for length biases during sequencing ^59^. During read processing (see below), deduplication was not performed on the spike-in sequences due to their high abundance. We noted that the bias ratios of 18-mer to 30-mer remained about the same (between 1.47-1.67) between samples, and thus correcting for these ratios between experiments was not necessary. The spike-in sequences are listed in Table S1. When we controlled for their abundance, the trends in the length distribution plots (Figures 4C and 5D) remained. The ribosome profiling data was processed and signatures of altORF translation were assessed as before ^19^. Sequencing was performed on Hiseq3000, Novaseq 6000 or X (single end 50) at the National Heart, Lung and Blood Institute (NHLBI) DNA Sequencing Core or National Institute of Diabetes, Digestive and Kidney Diseases (NIDDK) Genomics Core.

#### Analysis of translation for altORF analysis

For analysis of ribosome profiling data, we implemented a reduced-transcriptome alignment (bowtie1 with -y option) and python analysis pipeline as described elsewhere ^19,60^. We used the Ref-Seq Select+MANE (ncbiRefSeqSelect), as downloaded from UCSC on 14 April 2020, for this alignment. We note that in the case of conventional 25-34 nt footprinting in Fig 3 and S3), we allowed 2 mismatches (-v 2 option in bowtie1). In datasets including short footprints (15-34 nt) (Figs 4-5 and S4-S5), we aligned the reads to the reduced transcriptome (as noted above but also with removal of duplicate entries using SeqKit software rmdup function. without allowing any mismatches to reduce noise from cases where reads could map to multiple sites (-v 0 option in bowtie1). As an additional measure to eliminate noise during short footprint mapping, prior to alignment removal of a single 5’ base was performed on all reads to eliminate any incorporated untemplated base during the reverse transcription step by using cutadapt2. Doing this eliminates the possibility that a read is rejected due to a mismatch from a 5’ untemplated base. Despite loss of this base, read sizes are still plotted according to their length before trimming.

We used our previously established custom code ^12,60^ to count reads and create plots of average footprint levels (Figs 3-5, S3-S5). We calculated rpm values for 5’ UTR, CDS and 3’UTR for each annotated reference sequence transcript. These values were then used to calculate 3’UTR:CDS and 5’UTR:CDS values (minimum raw reads > 5, genelist function). We also used these tools to create metagene average plots (Figure 3-5) using the (metagene and posavg functions). In general, we assume a shift of 12 nt between the 5’ end of a read and the P site of the ribosome, allowing us to plot approximate P sites in Figure 3 and S3. In contrast, Figs 4-5 and S4-S5 show data that is plotted based on assignment to the very 3’ end of the read. Distribution of the bowtie-mapped footprint sizes were calculated by fastqc (Babraham Bioinformatics) and plotted by Prism software (version 10.2.2). The 3’ di-nucleotide distribution for 15-18 nt (16-mer) footprints (Figure 4D and S4B) was created by custom python scripts (see Guydoshlab github, endmotif_distribution function) and was plotted by Prism software.

#### Analysis of poly(A) sequences in footprints

Ribosome profiling footprints generated by ribosomes that had partially moved into the poly(A) tail are not expected to align to the genome or transcriptome. To count these footprints, we used processed but not aligned reads to assess the number of ribosomes that are near the poly(A) tail on mRNAs (Figure 3E), similar to previous analysis^43^. We counted the number of consecutive As (8-18 nt) at the 3’ end of the footprints using a grep command in bash. Then we calculated the percentages of these reads in the total unmapped reads and plotted those with Prism software (10.2.2).

### Western blot analysis

To visualize differences in kinase phosphorylation levels and to monitor KD of Pelo in A549 cells, 10 µl of ribosome profiling lysate (see preparation above) was mixed with equivalent amount of SDS Sample Buffer (Invitrogen) then run on 4-20% gradient Mini-PROTEAN Tris-HCl gel (BioRad). 5-10 µl of samples prepared this way was run on a 4-20% gradient Mini-PROTEAN Tris-HCl gel (BioRad) or 7% PhosTag (Wako) gel prepared according to manufacturer’s instruction and as described before ^15^. Proteins were transferred to a 0.2 µm PVDF membrane using the Trans-Blot Turbo System (BioRad, TransBlot protocol for 4-20% gradient Mini-PROTEAN Tris-HCl gel or 1.3 A for 14 minutes for PhosTag gel) and membranes were blocked in EveryBlot Blocking Buffer (BioRad) for 10 minutes at room temperature. This was followed by incubation with primary antibodies overnight in TBST (Tris-buffered saline plus 0.1% tween) and 1% milk. The following antibodies were used: anti-p38 MAPK (Cell signaling Technologies #9212, 1:2000 dilution), anti-P-p38 MAPK (Cell signaling Technologies, 1:2000 dilution, #9211), anti-ZAK (Bethyl A301-993A, 1:2000), anti-P-JNK (Thr183/Tyr185) (Cell signaling Technologies #4668, 1:1000), anti-JNK (Cell signaling Technologies #9252, 1:1000),

Anti-H3 (Abcam #1791, 1:5000), anti-PELO (ThermoFisher, #PA5-31697, 1:1000). After washing three times for 5 minutes in TBST, secondary antibodies were incubated with the PVDF membrane for 1 hour at room temperature (Goat anti-rabbit antibody, BioRad, 1:3000). Then, after additional washes (3 times, 5 minutes each) in TBST, the PVDF membrane were incubated with Clarity Western ECL substrate (BioRad) for 5 minutes and proteins were visualized by Amersham Imager 600. Experiments were performed at least three times using biological replicates and original raw data will be available on Mendeley Data. A third replicate (Figure 5B) using a different lysis procedure was also consistent. In this case cells were lysed in 10 mM Tris-HCl, pH=7.5, 150 mM NaCl and 1 % Triton-X on ice for 5 minutes and then centrifuged at 1500 rpm, for 4 minutes at 4°C. Then the supernatant was mixed with equivalent amounts of SDS sample buffer before loading it onto the gels.

